# Anatomical drivers of stomatal conductance in sorghum lines with different leaf widths grown under different temperatures

**DOI:** 10.1101/2022.10.16.512409

**Authors:** Yazen Al-Salman, Francisco Javier Cano, Ling Pan, Fiona Koller, Juan Piñeiro, David Jordan, Oula Ghannoum

## Abstract

Improvements in leaf water use efficiency (*iWUE*) can maintain crop productivity in water limited environments under rising temperatures. We investigated the leaf anatomical traits which underpin our recently identified link between leaf width (*LW*) and *iWUE*.
Ten sorghum lines with varying *LW* were grown under three temperatures to expand the range of variation of both *LW* and gas exchange rates. Leaf gas exchange, surface morphology and cross-sectional anatomy were measured and analysed using structural equations modelling.
Narrower leaves had lower stomatal conductance (*g_s_*) and higher *iWUE* across growth temperatures. They also had smaller intercellular airspaces, stomatal size, percentage of open stomatal aperture relative to maximum, hydraulic pathway, mesophyll thickness, and leaf mass per area. Structural modelling revealed a developmental association among leaf anatomical traits that underpinned *g*_s_ variation in sorghum.
Growing temperature and *LW* both impacted leaf gas exchange rates, but only *LW* directly impacted leaf anatomy. Wider leaves may be more productive under well-watered conditions, but consume more water for growth and development, which is detrimental under water stress.

**Highlight:** Coordination between leaf width and leaf anatomy underpins stomatal conductance variation in sorghum grown under different temperatures.

## INTRODUCTION

Growing population requires increased agricultural production per unit of arable land (Ray *et al*., 2013; Challinor *et al*., 2014; Valin *et al*., 2014). Since the majority of freshwater is utilised for agriculture (Postel *et al*., 1996), it is critical to breed new crop varieties in rain-fed agriculture that efficiently use available soil water (Condon *et al*., 2004; Blum, 2009). Climate change is expected to lead to increased air temperature, resulting in yield losses (Alexander *et al*., 2006; Lobell *et al*., 2013). Water use efficiency (*WUE*), also known as transpiration efficiency, expressed as the ratio of net carbon assimilation rate (*A_n_*) to transpiration rate (*E*), has proven to be an effective selection trait for crop improvement (Condon *et al*., 2004; Blum, 2005). *E* is determined as the product of the total leaf conductance to water vapour (including stomatal conductance (*g_s_*), cuticular conductance and the boundary layer conductance) with the leaf-to-air vapor pressure deficit (*VPD*), which depends on ambient air temperature and relative humidity, but also on leaf temperature (Jones, 2013; Grossiord *et al*., 2020). By contrast, intrinsic *WUE (iWUE*), defined as the ratio of *A_n_* to *g_s_*, accounts directly for variables controlled by the plant (Osmond *et al*., 1980). Both *g_s_* and *A_n_* are influenced by leaf anatomy and biochemistry as well as environmental conditions, and reflect better the genetic control on *WUE* (Chaves *et al*., 2016; Leakey *et al*., 2019).

Crops that utilise C_4_ photosynthesis are vital to the global food supply (Leakey, 2009). They dominate tropical and subtropical regions (Edwards *et al*., 2010), where agricultural yield is particularly vulnerable to climatic changes (Challinor *et al*., 2014; Jägermeyr *et al*., 2021; Watson-Lazowski and Ghannoum, 2021). Domestication and breeding led to extensive selection for higher yield in C_4_ crops and greater stability of photosynthesis (Duvick, 2005). However, the chances for solely manipulating carbon gain in C_4_ crops as a means of attaining higher *iWUE* remain low (Leakey *et al*., 2019). Alternatively, increasing *iWUE* in rain-fed C_4_ crops, such as sorghum, by reducing *g*s can potentially extend water availability until grain filling and limit the impact of water stress on yield (Ghannoum, 2016; Jackson *et al*., 2016; George-Jaeggli *et al*., 2017; Prasad *et al*., 2019). Hence, an emerging area of research in C_4_ crops is focused on reducing *g_s_* with minimal penalization in *A_n_* to enhance *iWUE*, following on similar advances in C_3_ plants (Yang *et al*., 2016; Bertolino *et al*., 2019; Leakey *et al*., 2019; Pathare *et al*., 2020; Israel *et al*., 2022; Pan *et al*., 2022).

Stomatal conductance is controlled by the number, dimensions and clustering of stomatal pores (Hetherington and Woodward, 2003; Franks and Beerling, 2009a; Lehmann and Or, 2015; Harrison *et al*., 2020). Stomata in grasses form paired dumbbell-shaped guard cells, flanked by two subsidiary cells. The linear shape of dumbbell guard cells, and smaller volumes, requires a smaller change in turgor pressure to generate the same pore aperture, allowing for faster stomatal movements than the kidney-shaped guard cells of many eudicots (Hetherington and Woodward, 2003; Franks and Farquhar, 2007; Harrison *et al*., 2020). Leaves with higher *g_s_* usually have higher stomatal density (*SD*) and smaller stomatal size (*SS*) compared to lower *g*_s_ leaves (Franks and Beerling, 2009b, 2009a). Smaller stomata occupy less epidermal space and reduce the distance traversed by gas molecules through the pore (Franks and Farquhar, 2007; Franks and Beerling, 2009a). Moreover, smaller guard cells have higher surface area in contact with neighbouring cells by unit guard cell volume, facilitating the efficient exchange of ions and osmolytes, hence requiring a smaller change in turgor when changing pore size (Dow *et al*., 2014; Lawson and Blatt, 2014). Higher *SD* and vein density (*VD*) reduce the hydraulic pathway from the xylem to the stomata (*Dist_H_*), promoting more efficient hydraulic supply but enhancing the risk of tissue dehydration (Fiorin *et al*., 2016; Brodribb *et al*., 2017). Reduced *SS* in plants developing under water stress is usually linked with higher *iWUE* (Franks and Farquhar, 2001; Doheny-Adams *et al*., 2012; Zhao *et al*., 2015). Also, *SS* has been genetically linked to *iWUE* in *Arabidopsis* (Dittberner *et al*., 2018). Therefore, adjustments in *SD* and *SS* may influence *iWUE*, although the combinations of *SS* and *SD* that improve *iWUE* seem species specific (Hughes *et al*., 2017; Bertolino *et al*., 2019; Caine *et al*., 2019; Dunn *et al*., 2019). The active control of guard cells aperture also significantly contributes to *g_s_*, especially in graminoids (Franks and Farquhar, 2007).

In C_4_ grasses, stomatal development progresses axially starting at the leaf base, with guard and subsidiary cells forming in files adjacent to the longitudinal, parallel veins (Rudall *et al*., 2017; Hepworth *et al*., 2018; Baird *et al*., 2021). The number and position of stomata are also determined at the early stages of leaf development (Vatén and Bergmann, 2012; McKown and Bergmann, 2020). Hence, changes in the interveinal distance between longitudinal veins (*IVD*) can influence the distribution and frequency of stomata (Ueno *et al*., 2006; Way, 2012; Xiong *et al*., 2017; Reeves *et al*., 2018; Schuler *et al*., 2018; Pathare *et al*., 2020; Pan *et al*., 2022). *IVD* is also an important determinant of C_4_ photosynthesis, through its effect on the size and spacing of the photosynthetic tissues in C_4_ leaves (mesophyll (M) and bundle sheath (BS) cells), which develop from the meristem around the vasculature (Dengler *et al*., 1985, 1986, 1994; Langdale *et al*., 1989; Dengler and Nelson, 1999; Ogle, 2003; Mckown and Dengler, 2007; Rizal *et al*., 2015). Furthermore, the ‘one-cell spacing rule’, which states that stomata are separated from each other by at least one epidermis cell to ensure efficient functioning, implies that producing more or bigger stomata would increase the substomatal airspaces and hence the area for gas exchange by diffusion (Dow *et al*., 2014; Franks and Casson, 2014; Harrison *et al*., 2020; McKown and Bergmann, 2020; Pathare *et al*., 2020). Hence, the development of stomata, veins and photosynthetic tissues is inherently interlinked in C_4_ grasses, as previously observed across C3 species (Brodribb and Feild, 2010; Brodribb *et al*., 2013; Buckley *et al*., 2015), and this coordination likely influences *iWUE* in C_4_ crops.

Recently, Cano *et al*. (2019) found that leaf width (*LW*) correlated negatively with *iWUE* in diverse C_4_ grasses, due to a positive association between *LW* and *g_s_*. They hypothesized that wider leaves operate at higher *g_s_* to offset the impact of the thicker boundary layer (BL) on leaf temperature, as higher transpiration rate is required to cool down the leaf (Gates, 1968; Jarvis and McNaughton, 1986; Schuepp, 1993; Leigh *et al*., 2017). Reliance on higher *g_s_* to control leaf temperature in wider sorghum leaves was confirmed under field conditions (Pan *et al*., 2022). Pan *et al*. (2022) also illustrated the positive association between *LW*, *IVD* and the percentage of open stomatal aperture relative to maximum (*% aperture*), and the negative association of *LW* with *SD* and *iWUE*. However, this was a field-based study conducted under hot and moderate drought conditions likely masking the full impact of *LW* on *iWUE*. Furthermore, new studies are needed to understand how leaf anatomy influences *iWUE* in C_4_ crops and how *LW* modifies them. This could bring about the use of *LW* as a surrogate for *iWUE* in breeding practices in C_4_ crops as both *LW* and stomatal traits are highly heritable traits in sorghum (Liang *et al*., 1973, 1975; Zhi *et al*., 2022).

Temperature regulates biochemical and biophysical processes that in turn regulate leaf expansion and development (Ben-Haj-Salah and Tardieu, 1995; Lafarge *et al*., 1998), and C_4_ gas exchange (Sage and Kubien, 2007; Yamori *et al*., 2014). In fact, *LW* is positively correlated with growing season temperature among C_4_ grasses worldwide (Baird *et al*., 2021). Also, the relationships between *LW*, leaf anatomy and gas exchange at different growing temperatures remains largely unexplored in C_4_ species (Kemp and Cunningham, 1981; Stamp *et al*., 1984, 1985). Hence, we explored the impact of *LW* on leaf anatomy and gas exchange in sorghum lines with varying *LW* growing in well-watered conditions under different temperatures to test the following hypotheses: 1) *LW*, *A_n_* and *g_s_* will increase and *iWUE* will decrease with growth temperature; 2) narrower leaves will have lower *g_s_* and higher *iWUE* across growing temperatures; 3) narrower leaves will have higher *SD* and *VD*, leading to shorter hydraulic path (*DistH*) and smaller intercellular airspaces (*IAS);* 4) shorter *DistH* and smaller *IAS* will correlate with smaller *SS* and stomatal apertures leading to lower *gs* and higher *iWUE*.

## MATERIALS & METHODS

### Plant material

Ten *Sorghum bicolor* (L. Moench) lines (**Table S1**) were randomly selected from more than 500 accessions of the sorghum conversion program (SCP) to meet two criteria: low tillering and a wide range of *LW* without any previous knowledge of *WUE*. The SCP is a backcross breeding scheme in which genomic regions conferring early maturity and dwarfing from an elite donor were introgressed into approximately 800 exotic sorghum accessions representing the breadth of genetic diversity in sorghum (Stephens *et al*., 1967). Our analysis was done at the level of line which included three replicates.

### Plant culture & experimental design

Seeds were germinated on 10^th^ of October 2016 in trays of 3 cm depth filled with soil and kept under controlled conditions of 25 °C, 60% relative humidity in darkness for 24 h and then moved inside a naturally-lit glasshouse at 22°C. Four days after germination plants were initially transplanted into 10 cm long pots and moved to their respective temperature treatments in three adjacent chambers described below. After one week, plants were transplanted into 7.5 L cylindrical pots of 40 cm depth to allow development of a deep root system. The soil substrate used throughout the experiment was a blend of soil, sand and organic material such as decomposed bark (Turtle Nursery, Windsor, NSW, Australia). The particle size promoted good drainage and aeration, and avoided plant roots drowning in the pots. 18.5 g of slow-release fertilizer (Osmocote^®^ Plus Organic All Purpose) with N-P-K of 13.4-1.0-5.2 (%) was added per pot. Plants were watered regularly to ensure no water stress.

Three temperature treatments were implemented in three adjacent chambers (8m long x 3m wide x 5m tall) in a naturally-lit, controlled environment glasshouse (Plexiglas^®^ Alltop SDP 16; Evonik Performance Materials, Darmstadt, Germany) at the Hawkesbury Institute for the Environment, Australia (−33.612032, 150.749098). The temperature treatments were designed to cover the range of average summer temperatures experienced by sorghum crops in its different areas of production (Ciampitti *et al*., 2019). The mean temperatures during the light period were 22.3°C, 28.7°C and 34.6°C, respectively for the three treatments (22, 28 and 35 °C hereafter) (**Fig. S1a**). A typical diurnal range of ~11°C was maintained in all treatments by heating and cooling throughout the day-night cycle. Relative humidity was kept close to 60% in the three glasshouse chambers (Carel Humidisk 65 humidifier), leading to maximum VPD of 0.9, 1.3 and 2.6 kPa for each temperature treatment (**Fig. S1b**). The photosynthetic photon flux density (PPFD) at canopy height was measured with a quantum sensor (Apogee quantum sensor, USA), and varied with prevailing weather conditions but was equivalent across rooms often reaching maximum PPFD ~1500 μmol m^-2^ s^-1^. Pots were randomly distributed within the glasshouse and rotated every week within each chamber and every two weeks between chambers to minimize chamber effect.

### Leaf gas exchange

Leaf level gas exchange was measured using LI-6400XT infra-red gas analysers (LI-COR Biosciences, Lincoln, Nebraska). The 8^th^, 9^th^ or 10^th^ fully expanded (depending on the plant growth rate) and mature leaf, usually two weeks after ligule was visible, was selected for measurement. Steady state gas exchange measurements were done in the flatted region of the leaf lamina, avoiding midrib, 40-45 days after emergence (DAE) when plants were at the vegetative stage and before head formation. Conditions inside the LI-6400XT chamber were 400 μmol CO_2_ mol^-1^, PPFD of 2000 μmol photons m^-2^ s^-1^ (10% blue light), and the block temperature was set to match the respective growth temperature (leaf temperature, measured with the internal thermocouple, reached 28°C, 35°C and 41 °C for the 22, 28 and 35 treatments, respectively). Gas exchange was measured only during sunny days at midday. Main variables extracted were net carbon assimilation rate (*A_n_*), stomatal conductance (*g_s_*) and intrinsic water use efficiency, *iWUE* (*A_n_*/*g_s_*).

### Leaf anatomy and leaf mass per area

A 1 cm long section spanning the width of the leaf (1 cm **x** *LW*), was cut at the middle of the leaf, at the same position previously used for gas exchange measurements two days before. Leaf sections were fixed in a mixture of formaldehyde, acetic acid and 70% ethanol in 5: 5: 90 proportions respectively for 24h. Subsequently they were kept in 70% ethanol and in darkness at room temperature. This section was used for stomatal, vein and inner leaf anatomy measurements. Below this 1 cm section, 4 leaf discs (1 cm^2^ each) were sampled and oven dried at 70 °C for 48 hrs. The dried leaf material was weighed. Leaf mass (g) was divided by disc area (m^2^) to calculate leaf mass per area (*LMA*, g m^-2^).

### Stomatal traits

For stomatal measurements, two small leaf sections between the 2^nd^ and 3^rd^ major veins were cut, one from each side of the midrib. Confocal microscopy (Leica TCS SP5, Leica Microsystems) was used to image the adaxial and abaxial surfaces of both sections, producing 4 images per leaf at x10 magnification. Images from the microscope were analysed using Image J (Schneider *et al*., 2012). Stomatal density (*SD*) was calculated as the number of stomata per unit area of the four images and averaged by leaf area (*SD* = *SD*_adax_ + *SD*_abax_). Within each area where *SD* was calculated, ten stomata were randomly selected to measure (**Fig. S3**): stomatal complex width (*W_s_*, including two guard cells and two subsidiary cells) and guard cell length (*L_s_*) to calculate stomatal size (*SS* = *W_s_* x *L_s_*), expressed in μm^2^; Epidermal cell size (*ES*) was calculated by dividing the area of the epidermal layer (minus the stomatal area) by the number of epidermal cells in that area, expressed in μm^2^. Maximum pore aperture (*a_max_* in μm^2^) was calculated as:

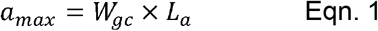

where *W_gc_* = width of the two guard cells of closed stomata and *L_a_* = is the length of the stomatal pore. This formulation was used because the shape of a fully open stomatal pore geometrically fits a rectangular shape in grasses (Franks and Farquhar, 2007; Franks *et al*., 2014). The fraction of the epidermis allocated to stomata (*f*_gc_) was calculated per de Boer *et al*. (2016), by dividing *SD/2*, to account only for one leaf side, times *SS* (in mm^2^) and expressed in % as follows:

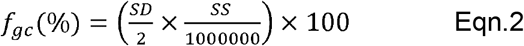

Theoretical maximum conductance (*g_smax_*) was calculated following Franks and Farquhar (2001), which includes one end correction:

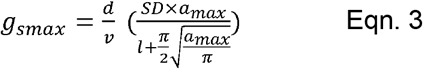

where *d* = diffusivity of water vapor in air; *v* = molar volume of air; *l* = stomatal pore depth which is assumed to be equivalent to *W_gc_* / 2. *d* and *v* were corrected for atmospheric pressure and temperature at each growing treatment (**Methods S1**). Finally, the operational stomatal pore (*a_op_*) that matched the corresponding measured *g_s_* was calculated following Pan *et al*. (2022):

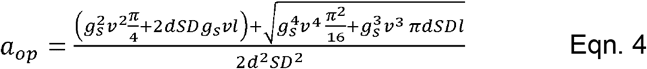

The ratio of the operational stomatal pore area (*aop*) to the maximum pore aperture (*a_max_*), expressed as a percentage, was termed % *aperture*.

### Vein traits

A 5 mm long section from the same 1 cm leaf section fixed earlier was cut and put through a clearing and staining protocol outlined by Scoffoni and Sack (2013). After mounting the cleared and stained leaf section on slides, leaves were imaged under x10 magnification using a light microscope (Axio Scope. A1, Carl Zeiss Microscopy GmbH, Jena, Germany), and images were analysed using image J (Schneider *et al*., 2012). Analysis was carried out on the area of the leaf between the 2^nd^ and 3^rd^ major veins, where stomatal traits were calculated. The inter-veinal distance (*IVD* in μm) was defined as the average distance between each two minor longitudinal veins (**Fig. S4**). Vein density (*VD*) was calculated as vein length per area (mm mm^-2^), including both longitudinal and transverse veins.

### Leaf cross-sectional anatomy

From the same 1 cm leaf section that was fixed, a cross section of the leaf was hand cut using two razor blades clumped together and spanning the leaf width. Leaf sections were cleared as mentioned above and rinsed with water to be mounted on a microscope slide and imaged under x10 magnification using the same light microscope. The same region between the 2^nd^ and 3^rd^ major veins was selected for analysis, and up to 3 images were taken for analysis. In each image, 2 separate regions were identified, each spanning from the middle of the vascular bundle of one vein to the middle of the vascular bundle of the neighbouring vein, with the combination of these two regions comprising two full veins (**Fig. S5**). In each region, total cross-sectional area of mesophyll (M), bundle sheath (BS), vascular bundle (VB) and both epidermal layers (including bulliform cells) were measured. Intercellular airspace (IAS) was calculated by subtracting the sum of the areas of the different cell types from the total area of the cross-section. Interveinal distance from leaf cross-sections (*IVD_c_*) was calculated as the distance between VBs. M cell length (*MC_length_*) was the average of the vertical and horizontal mesophyll cell lengths. Leaf thickness (*LT*) was measured at different several points within the region and mesophyll thickness (*MT*) compromised the inner distance between both epidermis at the minor vein. Single cell cross-sectional area of M (*MC_area_*) and BS (*BSC_area_*) cells were calculated by dividing total M or BS area by the number of M or BS cells. The ratio of M cross-sectional area to BS was called *M:BS*. The cross-sectional surfaces areas were standardized by dividing the total cross-sectional area of M, BS, IAS and VB by the corresponding *IVD_c_* to account for the variation of *IVD* within lines (*M_si_, BS_si_, IAS_si_, VB_si_* respectively). Finally, measurements of variables were averaged for the two regions in each image, and the resulting values from the three images were averaged to get the mean per leaf.

The hydraulic path that water travels from the *VB* to the sites of evaporation at the sub-stomatal cavity, or hydraulic distance (*Dist_H_*), was estimated following Brodribb *et al*. (2007) as the Euclidean distance of the hypotenuse of a rectangular triangle with one leg being the horizontal distance between the stomata and the middle of the nearest projected vein (*Dist_s-v_*), measured from paradermal leaf sections (**Fig. S4**), and the other leg was estimated as *MT* / 2:

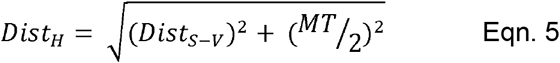

### Statistical Analyses

Statistical analysis and plotting were conducted using R software (https://www.R-project.org/). Normality was checked by fitting a generalized linear model and inspecting residual plots. Analysis of variance was carried out via a linear mixed-effects model (package “lme4”), with line as a random effect for within treatment analysis, and replicate as a random effect for within line analysis. In all analyses, variance within groups was performed using a *post hoc* Tukey test. The model for variation between temperature treatments included leaf width (*LW*) as a covariate effect and an interaction term as the fixed factors, while line was considered a random variable. The results of this analysis are presented in **Table 1**. A Pearson product-moment correlation analysis was performed to test statistical significance of relationships and obtain correlation coefficients (**Table 2**). To evaluate the relative contribution of different anatomical determinants on *g_s_*, a hierarchical partitioning analysis was ran using the package <u>hier.part</u>.

**Table 1.** Statistical output from a full-factorial mixed effects ANCOVA with leaf width as the covariate. This table also presents all the parameters measured with their abbreviations (*n*=3).

| Variables | df | Temperature | Leaf Width | Temperature * Leaf Width |
| --- | --- | --- | --- | --- |
| Net carbon assimilation rate ( $A_n$ ) | 1,2 | **<0.0001** | **0.015** | 0.6728 |
| Stomatal conductance ( $g_s$ ) | 1,2 | **<0.0001** | **<0.0001** | 0.6951 |
| Intrinsic water use efficiency ( $iWUE$ ) | 1,2 | **<0.0001** | **<0.0001** | 0.8761 |
| Leaf mass per area ( $LMA$ ) | 1,2 | **<0.0001** | **<0.0001** | 0.965 |
| Leaf width ( $LW$ ) | 1,2 | **<0.0001** | - | - |
| Stomatal density ( $SD$ ) | 1,2 | 0.9722 | 0.0556 | 0.6106 |
| Stomatal size ( $SS$ ) | 1,2 | 0.3408 | **<0.0001** | 0.1193 |
| Calculated maximum stomatal pore aperture ( $a_{max}$ ) | 1,2 | 0.0791 | **<0.0001** | 0.1753 |
| Epidermal cell area ( $ES$ ) | 1,2 | 0.7261 | 0.0557 | 0.335 |
| Operational stomatal aperture ( $a_{op}$ ) | 1,2 | **<0.0001** | **<0.0001** | 0.1067 |
| Percentage ratio of operational stomatal aperture to maximum pore size ( $\% aperture$ ) | 1,2 | **<0.0001** | **0.0013** | 0.6903 |
| fraction of leaf epidermis allocated to stomata ( $f_{gc}$ ) | 1,2 | 0.6247 | **0.0267** | 0.8017 |
| Theoretical maximum stomatal conductance ( $g_{smax}$ ) | 1,2 | 0.2315 | 0.8749 | 0.8904 |
| Interveinal distance measured from leaf lateral sections ( $IVD$ ) | 1,2 | **0.0227** | 0.0725 | 0.1963 |
| Vein density ( $VD$ ) | 1,2 | 0.0407 | 0.1175 | 0.216 |
| Total number of longitudinal veins ( $TLV$ ) | 1,2 | **<0.0001** | **<0.0001** | 0.3525 |
| Leaf mesophyll thickness ( $LT$ ) | 1,2 | **0.0045** | **0.0002** | 0.4272 |
| Mesophyll to bundle sheath ratio ( $M:BS$ ) | 1,2 | 0.3882 | **0.0163** | 0.6215 |
| Mesophyll surface area per $IVD_c$ ( $M_{Si}$ ) | 1,2 | 0.1531 | 0.4433 | 0.9278 |
| Bundle sheath surface area per $IVD_c$ ( $BS_{Si}$ ) | 1,2 | 0.1526 | **<0.0001** | 0.8784 |
| Vascular bundle surface area per $IVD_c$ ( $VB_{Si}$ ) | 1,2 | 0.1361 | **0.0309** | 0.4685 |
| Intercellular airspace surface area per $IVD_c$ ( $IAS_{Si}$ ) | 1,2 | **0.0023** | **0.0001** | 0.1924 |
| Average mesophyll cell length, vertical or horizontal ( $MC_{length}$ ) | 1,2 | **0.0083** | 0.1298 | 0.3755 |
| Mesophyll cell area ( $MC_{area}$ ) | 1,2 | 0.0569 | 0.0366 | 0.3584 |
| Bundle sheath cell area ( $BSC_{area}$ ) | 1,2 | 0.0817 | **0.0001** | 0.4755 |
| Interveinal distance measured from leaf cross-sections ( $IVD_c$ ) | 1,2 | 0.2254 | **0.0008** | 0.0312 |
| Distance from the stomata to middle of nearest projected VB ( $Dist_{s-v}$ ) | 1,2 | **0.0005** | **0.0009** | 0.4551 |
| Hydraulic Distance ( $Dist_{H}$ ) | 1,2 | **0.0027** | **0.0001** | 0.2735 |

**Table 2.**
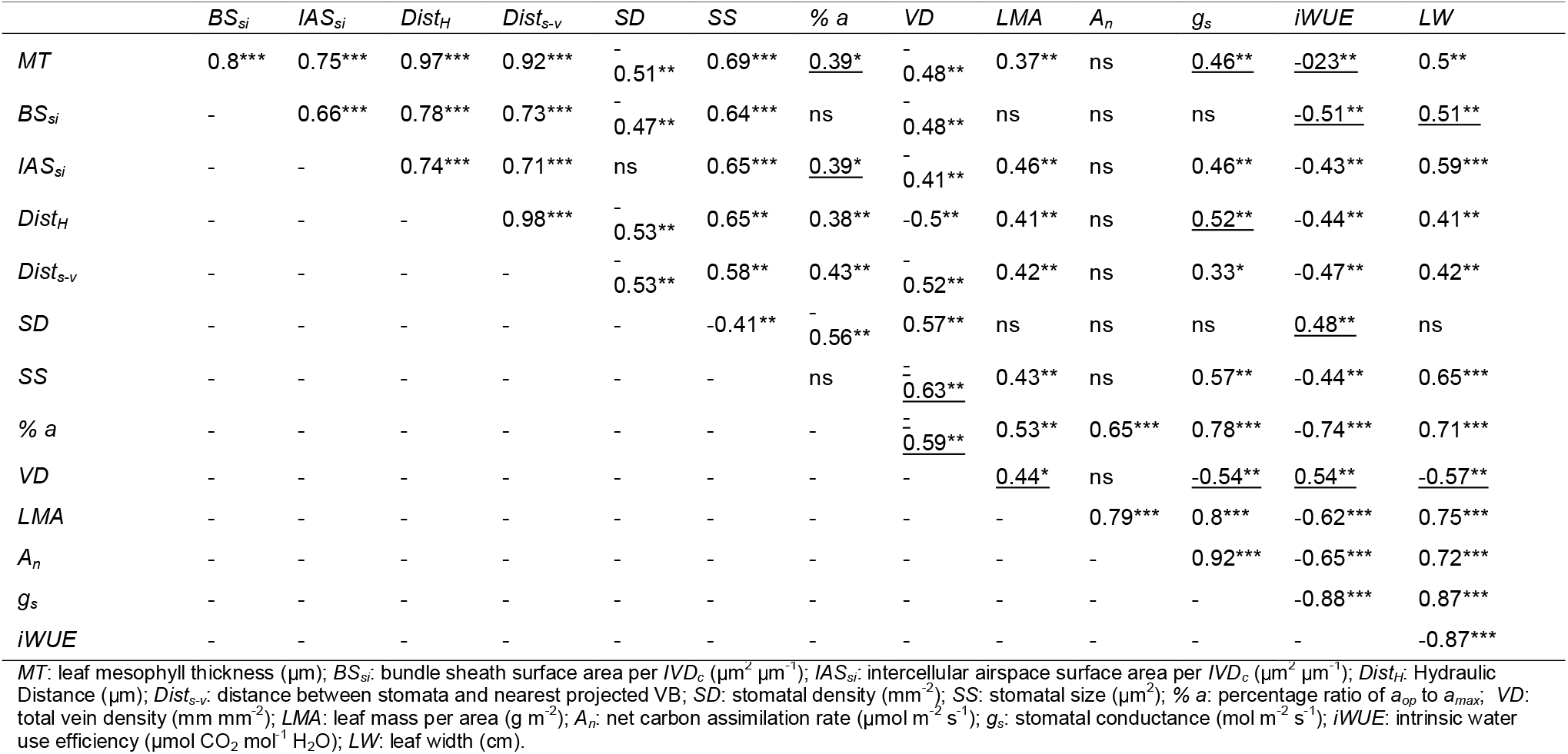
Pearson product-moment correlation analysis results for the relationships between the measured variables. The correlation coefficient and the statistical significance were determined using the mean value per line, per treatment for each variable. Statistical significance was judged as: *P*<0.001 (***), *P*<0.05 (**), *P*<01 (*), *P*>01 (ns). Underlined coefficients show the correlation between the variables was significant after excluding the 22°C treatment. (*n*=3).

To test and evaluate the multivariate and causal relationships underlying the observed effects of temperature on gas exchange variables via shifts in leaf anatomy and morphology we used structural equation modeling (SEM; Grace, 2006). First, SEM tested the adequacy of a causal model encompassing a set of a priori hypotheses which were established based on observed relationships among the drivers of *gs*, and hence *iWUE* (**Figs. 3** and **4**). We adopted Shipley’s d-separation method for model evaluation using the <u>piecewiseSEM</u> package in R (Lefcheck, 2016). This approach constructs SEM paths as a set of hierarchical linear models and has been suggested to have higher statistical power in studies with small sample size (Lefcheck, 2016; Shipley, 2016). A path coefficient is analogous to the partial correlation coefficient, and describes the strength and sign of the relationship between two variables (Grace, 2006). Mixed effect models were fitted using the “lme” function in the “nlme” package, while model assumptions were verified by inspecting residuals versus fitted values and quantile–quantile plots. We ran two analyses. First, we did not include temperature treatment in the SEM procedure, and we fitted linear mixed models for *g*_s_ and *A_n_* incorporating *LW* and a composite variable (i.e. *Anatomy*) as fixed effects, and line nested within temperature level as random factors. Further, we run a similar model including temperature as a predictor of *LW*, *Anatomy*, *g*_s_ and *A_n_* with line included as random factor as described previously, and temperature as fixed effect term in each of the remaining paths. The use of composite variables collapses the effects of multiple conceptually related variables into a single composite effect, simplifying the interpretation of model results (Grace, 2006). In our case we created the composite variable *Anatomy* using the variables that explained most of *g*_s_ variation: % *aperture, SS, IAS*_si_ and *Dist_H_*. Overall goodness-of-fit of the models was tested using Fisher’s C statistic (Lefcheck, 2016; Shipley, 2016). Non-significant *P*-values associated with goodness-of fit tests indicate acceptable model fit.

## RESULTS

### Genotypic variation and the effect of growth temperature on *LW* and gas exchange

Higher growth temperatures led to an increase in *LW* in most lines (**Fig. 1d**, **Table S1**; *P*<0.05). FF_SC842-14E had the lowest *LW* at each temperature (2.4 – 3.4 cm), while FF_SC1201-6-3 had the highest or second highest *LW* at the different temperatures (4.6 – 8.1 cm) (**Table S1**). *A_n_* and *g_s_* also increased with temperature (**Fig. 1a,b**). Within-line variation in *A_n_* was only apparent at 28°C and 35°C (**Table S1**). At the highest temperature, QL12, FF_SC56-14E and FF_SC1291-6-3 had the highest *A_n_* (> 50.5 μmol m^-2^ s^-1^), while FF_SC906-14E and LR9198 had the lowest (<45.0 μmol m^-2^ s^-1^) (**Table S1**). While lines changed rank based on mean *g_s_* at different temperatures, FF_SC842-14E maintained lowest *g_s_* (0.32 mol m^-2^ s^-1^), while FF_SC1201-6-3 had the highest (0.43 mol m^-2^ s^-1^). We found 15-20% variation in *iWUE* within lines (**Table S1**), but *iWUE* decreased with increasing temperature (**Fig. 1c**), confirming our first hypothesis.

**Fig. 1.**
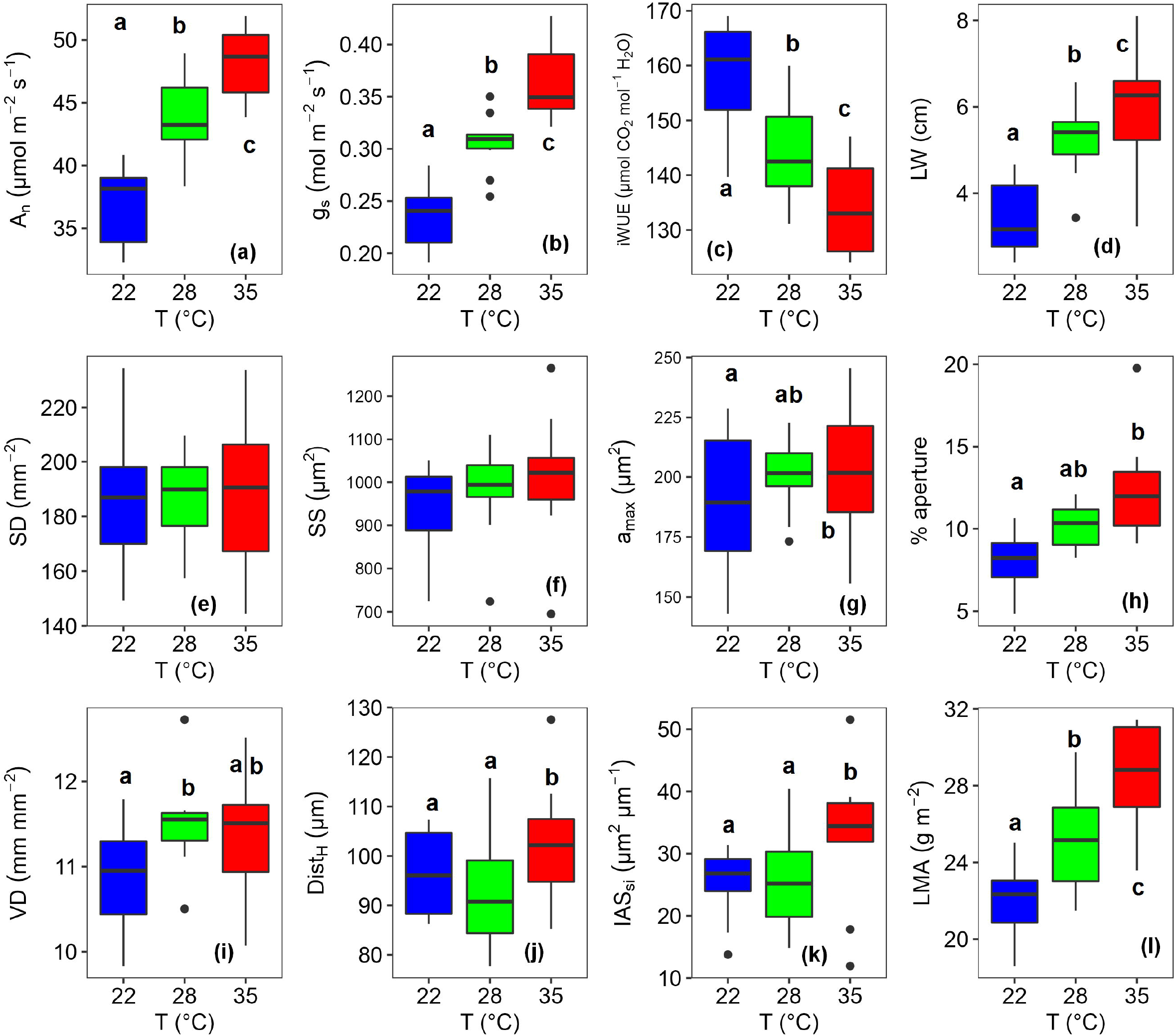
The distribution of trait values among the three temperature treatments. The distribution is summarized by boxplots for each treatment. The values in each boxplot compromise line mean for each treatment (n=3), yielding n=10 for each individual boxplot. Each box encompasses the 25th and 75th percentiles, with whiskers extending to show the extreme values of the measured variable. Statistically significant difference is represented at the top or bottom of each boxplot, with boxplots that share the same letters (a, b or c) having no significant difference between them, with significance taken at P<0.05. Different growth temperatures are represented by the different fill colour of the boxplot: blue=22°C, green=28°C, red=35°C. (a) Net carbon assimilation rate (*A_n_*); (b) Stomatal conductance (*g_s_*); (c) Intrinsic water use efficiency (*iWUE*); (d) Leaf width (*LW*); (e) Stomatal density (SD); (f) Stomatal size (SS); (g) maximum stomatal pore aperture (*a_max_*); (h) Operational stomatal aperture as percentage of maximum aperture (% *aperture*); (i) Vein density (VD); (j) Hydraulic distance (*DistH*); (k) Cross-sectional intercellular airspace surface area per *IVD* (*IASsi*); (I) Leaf mass per area (*LMA*).

### Correlations of leaf width with gas exchange

There was a positive relationship between *A_n_* and *g_s_* across temperatures (*R*=0.92, *P*<0.05; **Fig. 2a**). There were negative associations between *A_n_* and *iWUE* (*R*=-0.65, *P*<0.05; **Fig. 2c**), and between *gs* and *iWUE* (*R*=-0.88, *P*<0.05; **Fig. 2b**). This was accompanied by a strong positive correlation between *LW* and *g_s_*, (*R*=0.87, *P*<0.05; **Fig. 2e**), and a negative one between *LW* and *iWUE* (*R=-0.87, P*<0.05; **Fig. 2f**), even within growth temperatures (**Table S6**), confirming hypothesis 2. Within temperatures, *A_n_* did not correlate with *LW* or *iWUE*, but *g_s_* maintained that strong association (**Table S6**). Therefore, increases in *g_s_* with temperature were more influential than increases in *A_n_* on *iWUE*.

**Fig. 2.**
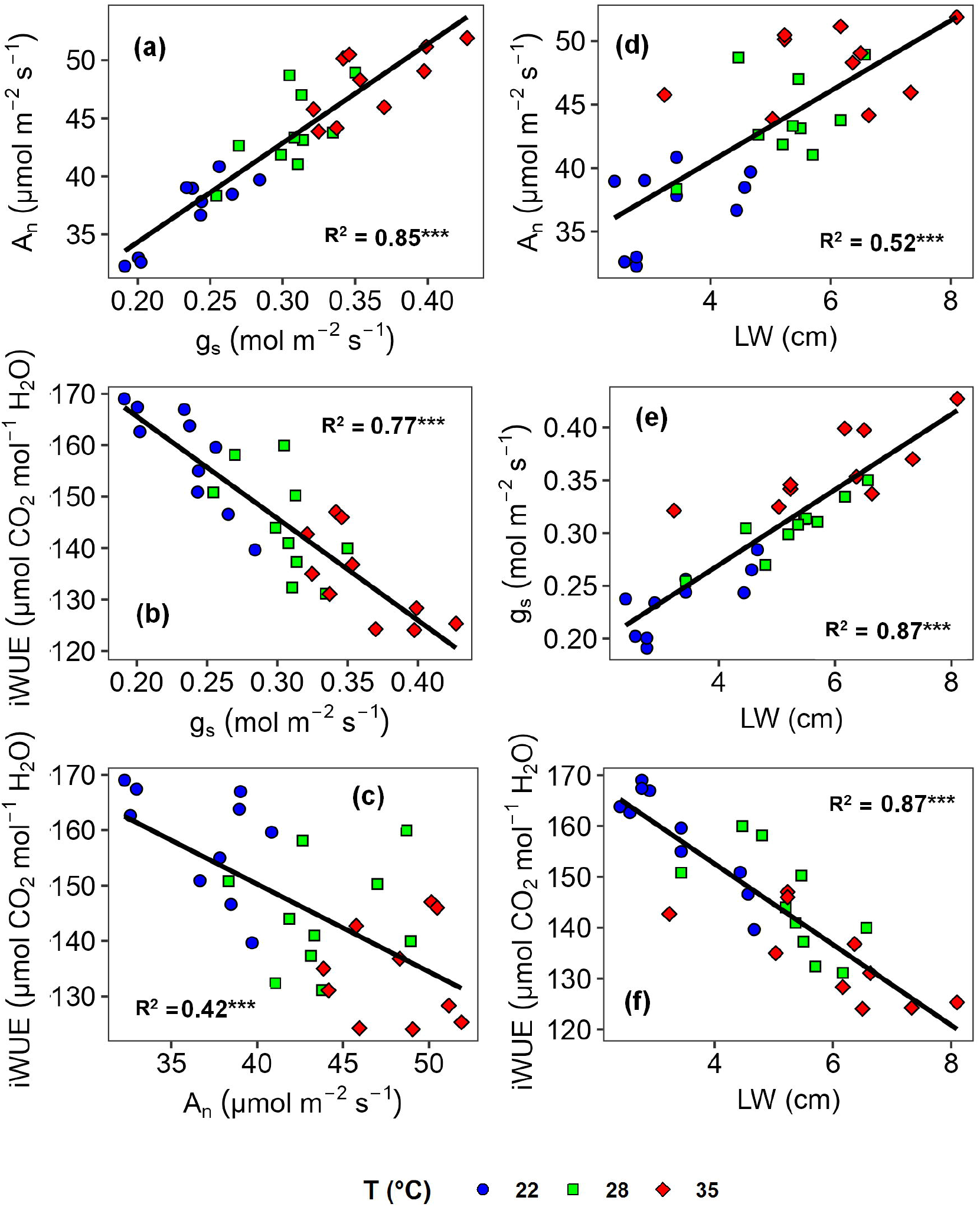
The relationship between leaf gas exchange parameters and leaf width under different growth temperatures in ten *Sorghum bicolor* lines. Data were collected on the youngest fully expanded leaf and measured at corresponding growth temperature and saturating light levels using the LI-6400XT. Each point represents the mean value of the variable per line and treatment (n=3). Standard error bars were removed to ensure clearer presentation (Table S1). Pearson correlation analyses were conducted and significant results were shown with a solid line with the corresponding *R^2^* value (P<0.001=***, P<0.05=**, P<01=*) (Table S5). Growth temperatures were: blue=22° C, green=28°C, red=35°C. (a) Carbon assimilation rate (*A_n_*) vs. Stomatal conductance (*g_s_*); (b) Intrinsic water use efficiency (*iWUE*) vs. *g_s_*; (c) *iWUE* vs. *A_n_*; (d) *A_n_* vs. Leaf width (*LW*); (e) *g_s_* vs. *LW;* (f) *iWUE* vs. *LW*.

### Anatomical determinants of *g_s_*

*g_s_* correlated positively with *SS* (*R*=0.57, *P*<0.05; **Fig. 3a**) and *a_max_* (**Table S5**), but did not correlate with *SD* (**Table 2**). However, there was a positive correlation between *g_s_* and theoretical maximum stomatal conductance (*g_smax_*) (**Table S5**), and between *g_smax_* and the fraction of epidermis occupied by stomata (*f_gc_*) (*R*=0.88, *P*<0.05; **Fig. 4a**). Anatomical variables correlating positively and significantly with *g_s_* included the percentage ratio of open stomatal aperture to its maximum aperture (% *aperture*) and cross-sectional surface area of intercellular airspaces (*IAS_si_*) (**Fig. 3b,c**). The correlations between *gs* and mesophyll thickness (*MT*), inter-veinal distance (*IVD*) and the leaf hydraulic distance (*DistH*) were was significant only after excluding the 22°C treatment from the global regression (**Table 2** and **Table S5**). However, the independent contribution of *Dist_H_* to *g*_s_ was higher (21%) than that of *MT* (14%), *IVD* (12%) and *SD* (4%), when % *aperture* was excluded and only strictly structural variables were considered (**Fig. 4d**). Furthermore, *g*_s_ was positively correlated with the projected distance from stomata to VB (*Dist_s-v_*) (**Table 2**). Other structural traits that were strong determinants of *g*_s_ according to hierarchical partitioning were *SS* (22%) and *IAS_si_* (25%) (**Fig. 4d**).

**Fig. 3.**
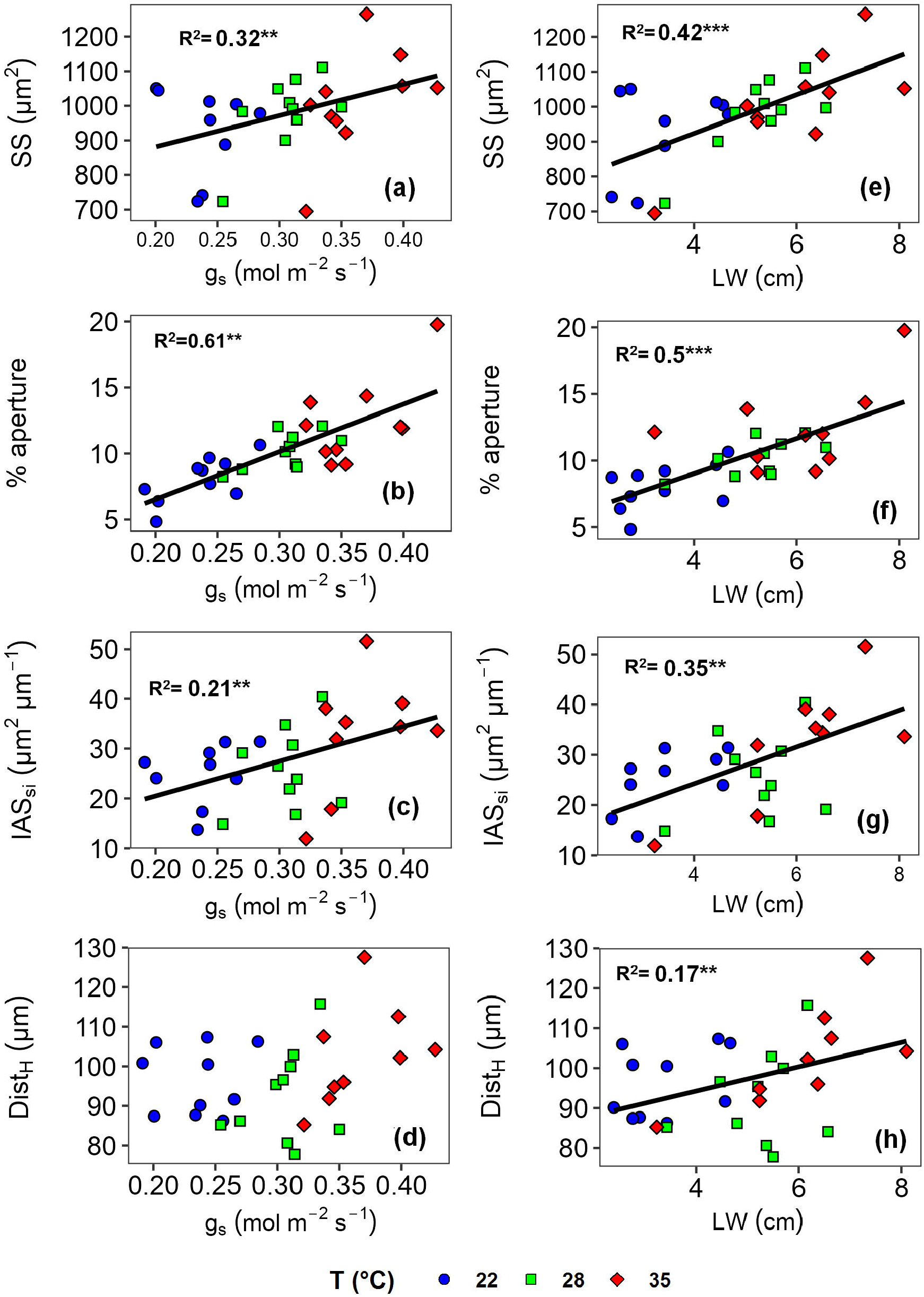
The relationship between leaf width (*LW*), stomatal conductance (*g_s_*) and anatomical traits linked with *g_s_* in ten *Sorghum bicolor* lines grown under different temperatures. Stomata, vein and inner anatomy traits were measured on the same portion sampled from the middle of the youngest fully expanded leaf. The area between 2^nd^ and 3^rd^ major veins from the midrib was analysed. Each point represents the mean per line and treatment (n=3). Standard error bars were removed to ensure clearer presentation (Tables S2, S3 and S4). Pearson correlation analyses were conducted and significant results were shown with a solid line with the corresponding *R^2^* value (P<0.001=***, P<0.05 =**, p<01=*). Different growth temperatures are represented by the different fill colour of the scatter points: blue=22°C, green=28°C, red=35°C. (a-d) Relationship between *g_s_* and (a) Stomatal size (SS), (b) operational stomatal aperture as percentage ratio of maximum aperture (*% aperture*), (c) Cross-sectional intercellular airspace surface area per *IVD* (*IAS_si_*), (d) hydraulic distance (*DistH*). (θ-h) Relationship between *LW* and (e) SS, (f) *% aperture*, (g) *IAS_si_*, (h) hydraulic distance *DistH*.

**Fig. 4.**
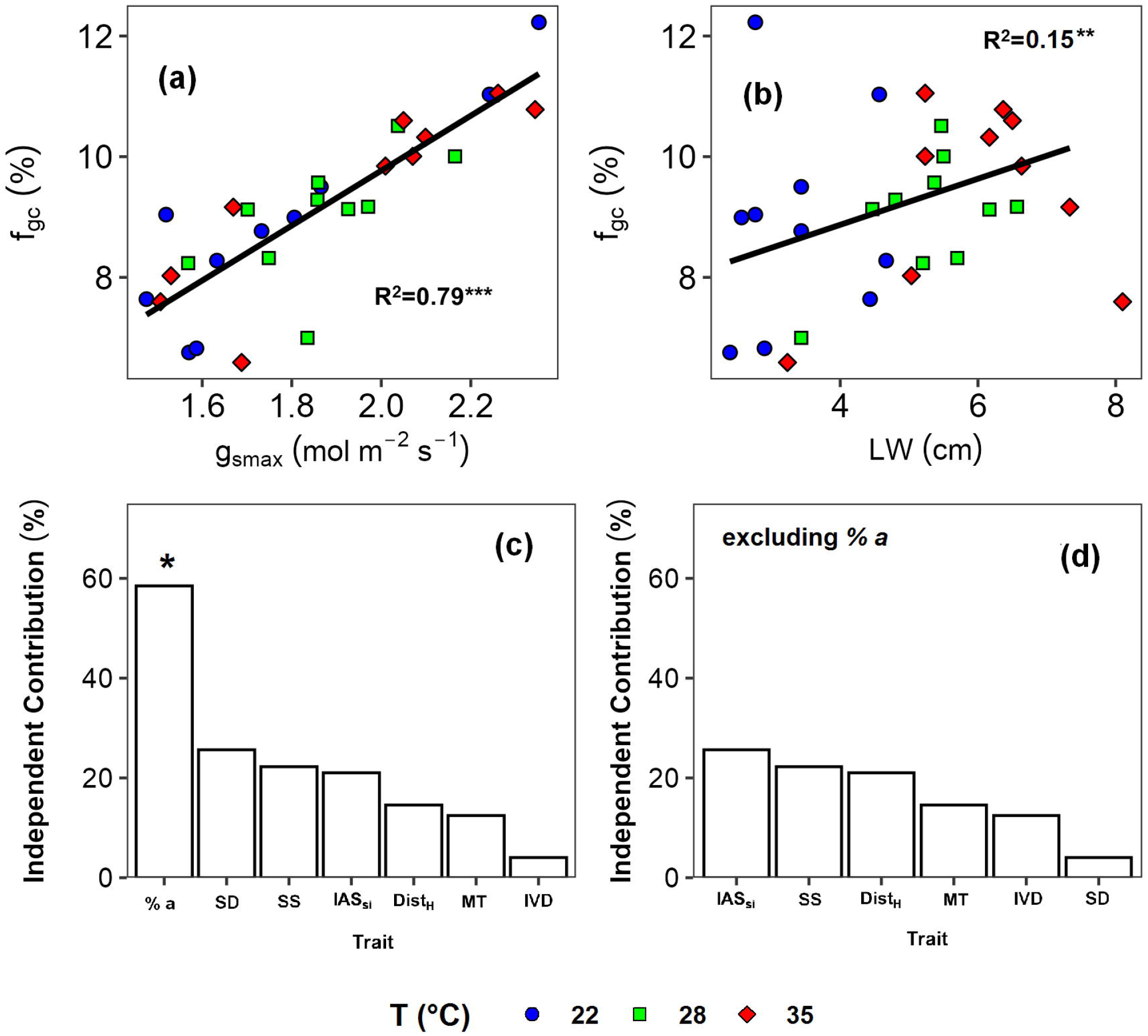
Anatomical determinants of stomatal conductance (*g_s_*) in sorghum. Data was collected on the youngest fully expanded leaf and measured at corresponding growth temperature and saturating light levels using the LI-6400XT. The same leaf was then sampled for anatomy with the area between 2nd and 3rd major veins at the middle of the leaf length was analysed for stomatal traits, (a) Percentage fraction of leaf epidermis allocated to stomata (*f_gc_*) vs theoretical maximum conductance (*g_smax_*); (b) *f_gc_* vs leaf width (*LW*); (c) variation by *g_s_* explained independently using a hierarchical partitioning analysis; (d) variation by *g_s_* explained independently using a hierarchical partitioning analysis excluding % *aperture* (% *a*). Significant contributions, indicated by an asterisk (P<0.05=*; see Materials & Methods), were determined afterwards using randomization tests. Abbreviations can be found in Table 1. Pearson correlation analyses in Figs, (a) and (b) were conducted and significant results were shown with a solid line with the corresponding *R^2^* value (P<0.001=***, P<0.05=**, P<01=*). *R^2^* value in Fig. b excludes the outlier point with *LW* higher than 8 cm (line FF_SCI 201-6-3). Different growth temperatures are represented by the different fill colour of the scatter points: blue=22°C, green=28°C, red=35°C.

### Correlation between leaf anatomical traits and leaf width

The *SS* was significantly correlated with other parameters that affected *g_s_*, namely *IAS_si_* and *Dist_H_* but was not correlated with *% aperture* (**Table 2**). *SD* (*P*<0.05; **Table 2**), cross-sectional bundle sheath surface area (*BS_si_*) and bundle sheath cell area (*BSC_area_*) (*P*<0.05; **Fig. S7b** and **S7d** respectively) were also significantly correlated with *SS*. *IAS_si_* and *BS_si_* also significantly correlated with *IVD* (*P*<0.05; **Table 2**). There was a strong correlation between *MT* and *Dist_s-v_* (*P*<0.05; **Table 2**).

*LW* correlated positively with *SS* (*R*=0.65, *P*<0.05), *% aperture* (*R*=0.71, *P*<0.05), *IAS_si_*, (*R*=0.59, *P*<0.05) and *DistH* (*R*=0.41, *P*<0.05) (**Fig. 4e,f,g,h** respectively). *LW* was also significantly and positively correlated with *fgc* (*R*=0.39, *P*<0.05; **Fig. 4b**), *g_smax_, LMA* and *MT* (*P*<0.05; **Table S5**), but there was no significant correlation between *LW* and *SD. LW* correlated negatively with *VD* only when considering the two higher growth temperatures (**Table 2**). Hence our third hypothesis was partially confirmed as we found strong connections among *LW* and several anatomical traits, but not with *SD*. In general, *LW* had a higher effect on leaf internal anatomy than temperature (**Table 1**).

### Network combination of traits

Structural equation modelling (SEM) was used to evaluate multivariate causal relationships among *LW*, leaf anatomy and leaf gas exchange. We fitted linear mixed models for the components of *iWUE* (*g*_s_ and *A_n_*) incorporating *LW* and a composite variable, *Anatomy*, that aggregated the main anatomical variables (*SS*, *IAS_si_*, % *aperture* and *DistH*) that were related to *LW* and *gs* (**Fig. 3** and **4**, **Table 2**). Our model combining growth temperatures and sorghum lines explained 84 % of *g_s_* variation, with both *LW* and *Anatomy* equally contributing to *g*_s_ variation (coefficient of 0.45 for both, *P*<0.001; **Fig. 5**), confirming hypothesis 4 that leaf anatomical traits shape *g*_s_. The strongest predictor of *A_n_* (78 %) was *g*_s_, but *LW* had direct positive control on *A_n_*, as well as an indirect positive effect mediated by its impact on leaf anatomy and *g*_s_ (**Fig. 5**). When temperature was explicitly considered into the SEM, temperature was not directly associated with *Anatomy*, but it was a strong determinant of *LW* and had independent positive effects on both *g*_s_ and *A_n_* of similar magnitude as the direct effects of *LW* and *Anatomy* on *g*_s_ (**Fig. S9**). However, once temperature was introduced, the strong positive connection between *LW* and *A_n_* found in the previous model became not significant, and *An* relied entirely on *g*_s_ and ambient temperature.

**Fig. 5.**
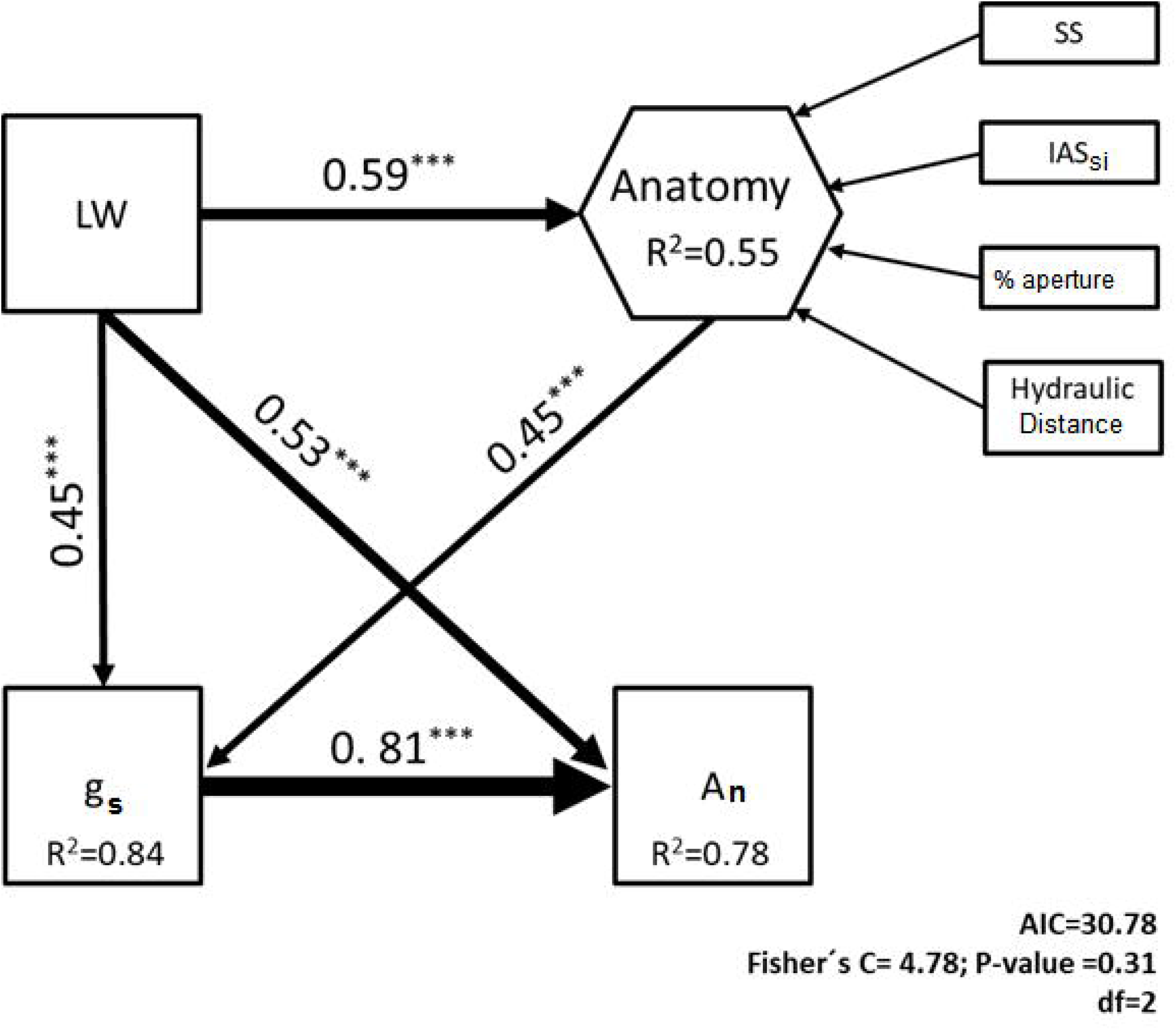
Structural equation analysis showing the effects of leaf width and leaf anatomy on stomatai conductance (*g_s_*) and carbon assimilation rate (*A_n_*) across temperature treatments and lines. Numbers adjacent to the arrows indicate the effect size of the relationship, while asterisks denote P-values at <0.0001. Leaf anatomy was included as a composite variable, and included the following variables: stomatai size (SS), Cross-sectional intercellular airspace surface area per inter-veinal distance (*IAS_si_*), operational stomatai aperture as a percentage of the maximum (% *aperture*) and hydraulic distance. *R^2^* denotes the proportion of the model variance explained. Overall goodness-of-fit test is shown in the bottom of the figure.

## DISCUSSION

### Wider leaves are associated with higher *g_s_* and reduced *iWUE*

Climate change is accelerating our need to improve the *WUE* of rainfed crops (Lobell *et al*., 2013). However, developing effective screening tools for *WUE* in C_4_ plants has been challenging (Ghannoum, 2016). Here, we showed that *LW* had strong positive influence on *g_s_* and a negative one on *iWUE* in sorghum under well-watered conditions (**Fig. 2e,f**). These results agree with the optimal size theory that states narrower leaves will be more water use efficient in hot and high-irradiance sites (Parkhurst and Loucks, 1972; Baird *et al*., 2021), as shown previously within C_4_ grasses (Cano *et al*., 2019; Pan *et al*., 2022) and in other plant species (Baldocchi *et al*., 1985). The negative effect of increasing *LW* on *WUE* was also recently observed across sorghum genotypes at the plant level (Zhi *et al*., 2022), where *WUE* (g kg^-1^) was determined as the ratio of total plant dry mass divided by total water use measured with lysimeters. Under hot and moderately dry conditions, the correlation between *LW* and *g*_s_ in sorghum was related to leaf temperature regulation (Pan *et al*., 2022), as wider leaves operated with higher *g*_s_, especially at low wind speed, to cool the leaf when ambient temperature was higher than the optimum for photosynthesis (Pan *et al*., 2022). Wider leaves have thicker boundary layers, which impose greater diffusive resistance to water vapour exiting the leaf through stomata, restricting convective heat transfer (Gates, 1968; Monteith and Unsworth, 2013). Hence, wider leaves get hotter more quickly than narrower ones under high radiation (Smith, 1978; Leigh *et al*., 2017). To counteract the greater boundary layer resistance, wider leaves increase *g*_s_ to drive transpiration and cool the leaf (Pan *et al*., 2022). The current study was conducted at a range of cool to warm temperatures, suggesting that factors other than temperature regulation control the positive association between *LW* and *g*_s_. Understanding this relationship is crucial due to the great agricultural interest in manipulating stomatal behaviour to increase *iWUE* (Jones, 2004; Lawson and Blatt, 2014; Leakey et al., 2019), in order to reduce the sensitivity of crops to drought stress associated with higher temperatures and vapor pressure deficits (Lobell *et al*., 2014).

### Modified stomatal anatomy in wider leaves supports higher *g_s_*

Consequently, how do wider leaves reach higher *g_s_* than narrower ones? There are three possibilities: 1) higher anatomical stomatal conductance (*g_smax_*), 2) more open stomata (increase *% aperture*), or 3) a combination of both. Our results corroborate the third explanation as we found evidence for both a *g_smax_*-driven increase in *g_s_* that is mainly associated with greater allocation of space to the stomata (**Fig. 4a**), as well as *% aperture-driven* increase in *g_s_* as the leaf widened (**Fig. 3f**). Indeed, the hierarchical model pointed out that *% aperture* was the most important variable that explained *g*_s_ when grouped with the strictly structural variables such as *SS* (**Fig. 4c**). Leaves with higher *g_s_* also had higher *% aperture* (**Fig. 3b**). Hence, active adjustment of the osmotic potential of guard cells was needed to increase their turgor pressure and support higher operational *g*_s_ irrespective of stomatal size and density (Hetherington and Woodward, 2003; Kollist *et al*., 2014; Israel *et al*., 2022). Increasing the operational stomatal aperture was observed in transgenic rice with reduced *SD* to achieve the same gas exchange rates as the wild-type rice in response to high irradiance and high temperature (Caine *et al*., 2019). Also, the rate of increasing osmotic concentration that opens stomata has been estimated to scale linearly with *SS*, assuming a constant flux rate of ions per unit area of guard cell plasmalemma (Raven, 2014). Due to *g_s_* also positively correlating with *LW*, *SS* and *g_smax_* in our study, sorghum lines with wider leaves may be considered energetically costly to operate compared with narrow leaves.

Furthermore, the cost-benefit analysis proposed by de Boer *et al*. (2016) relates the ‘benefit’ of having a higher *g_smax_*, associated with higher photosynthesis rates, with the ‘cost’ of allowing a higher *fgc* (**Fig. 4a**). Bigger and more stomata are more costly to develop and operate, and the likelihood of penetration by pathogenic fungi is increased (Lawson and McElwain, 2016). Surprisingly, in de Boer *et al*. (2016), amphistomatous monocots (mostly grasses) were characterized by decreasing *f*_gc_ as *g_smax_* increased. High *g_smax_* is usually linked with high productivity and resource-use when growing in a resource rich environment, but lower *g_smax_* with relatively low *fgc* may be more advantageous in resource poor habitats (Lawson and McElwain, 2016). Wild grasses have experienced little evolutionary pressure to increase leaf gas exchange rates, and so they follow a more conservative strategy. On average, they have tougher leaves with longer life spans (Onoda *et al*., 2011) and notably higher cell wall mass per unit leaf mass (Onoda *et al*., 2017). However, sorghum as a domesticated crop may have experienced selection towards higher productivity in which *LW*, owing to its strong relationship with *g_s_* and *A_n_*, may have played a central role (Milla *et al*., 2015; Preece *et al*., 2017; Fu *et al*., 2019). Hence, wide leaf sorghum may be more productive, but at the cost of higher resource consumption, particularly water. In addition, the higher cost of increasing *f_gc_* increases the reliance on osmotic adjustment to operate the stomata, which can be detrimental under water stress (Pan *et al*., 2022).

### Smaller *Dist_H_* and airspaces in narrow leaves contributed to decreasing *g_s_*

The anatomical trait that most explained *g*_s_ variation other than *SS* was the area of sub-stomatal airspaces (*IAS_si_*) (**Fig. 4d**). The one-cell spacing rule of stomatal distribution requires each stomata to be accompanied by large sub-stomatal cavity to allow for efficient diffusion of gases from and towards the mesophyll (Harrison *et al*., 2020). Greater *IAS_si_* leads to greater surface area for water evaporation within the leaf, which helps to reduce leaf temperature through latent heat transfer if there is also lower stomatal resistance and hence higher water vapour gradient between sites of evaporation and stomatal pore (Buckley *et al*., 2017; Cernusak *et al*., 2018). The optimum temperature for *An* in sorghum is close to 38°C (Sonawane *et al*., 2017) and hence, plants in our study had no special need to reduce their leaf temperature. Given that leaf anatomy does not change once the leaf is mature, our data suggest that the anatomy of wider leaves was set to withstand elevated temperatures even when developed at lower temperatures.

The anatomical coordination between *LW* and *IAS_si_* (and other tissues within the leaf) rests on how grass stomata develop and expand in files next to the longitudinal veins. In grasses, the earliest cell developmental stages are basally confined, and cells expand and differentiate longitudinally towards the tip of the leaf. Stomata develop from epidermal cell files adjacent to those which overlie longitudinal veins. The two guard cells with the two subsidiary cells are differentiated moving away from the leaf base, forming the stomatal complex and the rest of epidermal cells (Vatén and Bergmann, 2012; McKown and Bergmann, 2020). The molecular regulation of *SD* remains unknown, although it was suggested that a vein derived morphogen may guide the number of and spacing between stomatal files flanking the veins (Kamiya *et al*., 2003; McKown and Bergmann, 2020). Our data support this hypothesis as wider leaves had larger distances from stomata to the vein (*Dist_s-v_*), which was associated with lower *SD* and higher *SS* and *IVD* (**Table 2**). Thus, the unknown morphogen might be produced at higher concentration around the veins of narrow leaves, and that higher morphogen gradient induced higher *SD* and lower *SS*. This mechanism explains commonly observed relationships, such as the general positive association between *LW* and *IVD* in grasses (Sage, 2001; McKown and Dengler, 2009; Smillie *et al*., 2012; Griffiths *et al*., 2013; Ruwanthi Nayananjalee *et al*., 2017; Baird *et al*., 2021; Pan *et al*., 2022), or the coordination between vein and stomatal densities (Carins Murphy *et al*., 2012, 2016; Brodribb *et al*., 2013, 2017; Zhang *et al*., 2018).

We also found that increased *IVD* in sorghum was linked to thicker mesophylls (*MT*), leading to increased *IAS_si_* as leaves widened, especially at higher growth temperatures (**Fig. S8**, **Table S5**). Furthermore, the strong positive correlation between *MT* and *Dist_s-v_* meant that the distance water must travel from the xylem to the sites of evaporation (*Dist_H_*) was associated with *IAS*_si_ and *LW*. The coordination among IAS, mesophyll cells and stomata to facilitate efficient hydraulic transport has been illustrated in wheat (Lundgren *et al*., 2019) and *Populus tremula* (Aasamaa *et al*., 2001). Longer *Dist_H_* in wider leaves can increase hydraulic resistance in living cells, hindering hydraulic supply and increasing the risk of leaf dehydration at high transpiration rates and/or lower leaf water potentials (Sack and Frole, 2006; Sack and Holbrook, 2006; Brodribb *et al*., 2007; Buckley *et al*., 2015; Fiorin *et al*., 2016). On the other hand, narrower leaves kept the water source for transpiration at shorter distances, reducing leaf hydraulic resistance, and making leaf water status more reliant on active stomatal control. Narrow leaves also shortened the sink for photosynthetic products (lower *BS_si_*), ensuring more efficient nutrient transport, likely contributing to their higher *iWUE*. While reduced *SD* increased *iWUE* in C_3_ grasses (Hughes *et al*., 2017), in the more complex anatomy and physiology of C_4_ photosynthesis, increased *iWUE* was not linked to reduced *SD* (almost the opposite), but to reduced *IAS*_si_, *Dist_H_, SS* and *% aperture*, all positively correlated with *LW* and negatively with *iWUE* (**Table 2**).

### Changes in biomechanical traits associated with leaf width

Finally, developmental coordination among anatomical traits is important for the biomechanical aspects of sorghum leaves. Wider leaves had higher *IVD* and lower *VD*, reducing the structural integrity of tissue between the veins. However, wider leaves also showed higher *IAS_si_* and *MT*, helping them maintain biomechanical strength. In particular, veins (sclerified structures) provide the leaf with support in order to withstand forces without breaking or causing damage to the xylem (Read and Stokes, 2006; Onoda *et al*., 2008, 2011; He *et al*., 2019). Increased *MT* enables the maintenance of “flexural stiffness” (Onoda *et al*., 2008; He *et al*., 2019). Wider leaves also had higher *BS_si_* and *VB_si_*, likely leading to their higher leaf mass per area (LMA) (**Table 2**). *LMA* also increased with *IVD, MT* and *IAS_si_* (**Table 2**), but leaf density was unrelated to *LW*, suggesting that wider leaves compensated the more porous tissue in between the veins with a reinforced and denser tissue in the veins (Onoda *et al*., 2011), albeit at the higher energetic cost of producing more lignin (Sage, 2004).

## Conclusion

We found significant correlations among distant cells which obey a coordinated developmental scheme that ensured higher *gs* but lower *iWUE* in wider sorghum leaves. Four anatomical traits strongly explained the variation of *g*s: *% aperture, IAS*si, *SS* and *DistH*, all positively correlated with *LW*, and negatively with *iWUE*. These correlations bring on effective heat dissipation within the leaf, but as these correlations were sustained across all the growing temperatures, we conclude that *LW*, a highly heritable trait, exerts strong control on leaf anatomy to withstand elevated temperatures irrespective of current growing temperatures. We also found biomechanical trade-off that ensured leaf support as leaves widened. Wider leaves may be more productive under well-watered conditions, but at the cost of higher water consumption and cost to develop. They also rely more on osmotic adjustment to operate their larger stomata, which is detrimental under water stress. Ultimately, we found that *LW* is pivotal for the regulation of *gs* in sorghum, directly and indirectly through modifying leaf anatomy. *LW* can be a good proxy for *iWUE* in breeding C_4_ crops for rain-fed conditions.

## ACKNOWLEDGEMENTS

This research was funded by the ARC Centre of Excellence for Translational Photosynthesis (ARC CoETP; grant no. CE140100015. awarded to OG and DJ and an EMCR seed grant from the ARC CoETP awarded to FJC. FJC was also funded through the Spanish fellowship Juan de la Cierva-Incorporación (IJC2019-041435-I). YA was supported by a PhD scholarship jointly awarded by the ARC CoETP and the Hawkesbury Institute for the Environment, Western Sydney University. We would also like to acknowledge the contributions of the Australian Grains Research and Development Corporation and the Queensland Government to the development of germplasm and on-going funding of the Queensland sorghum breeding program.

## AUTHOR CONTRIBUTION

YA and FJC are joint first authors. FJC and OG designed the experiment based on an original idea of FJC. FJC grew the plants, undertook the gas exchange measurements with help from FK. YA measured anatomical traits with help from LP and analysed and plotted all the data. JP conducted the structural equation modelling and interpretation with help from YA and FJC. YA, FJC and OG drafted the manuscript and all authors contributed to the editing. OG coordinated the project execution.

## DATA AVAILABILITY

The data generated and analysed during this study are available from the corresponding authors by request. Raw data includes gas exchange and anatomical analyses.

## SUPPORTING INFORMATION

**Methods S1** Calculations of water diffusivity in air and molar volume of air.

**Table S1** Means of measured leaf gas exchange, mass per area and width for each *Sorghum* line at the corresponding treatment.

**Table S2** Means of measured leaf stomatal anatomy parameters for each *Sorghum* line at the corresponding treatment.

**Table S3** Means of measured leaf vein anatomy parameters for each *Sorghum* line at the corresponding treatment.

**Table S4** Means of measured inner leaf anatomical parameters for each *Sorghum* line at the corresponding treatment.

**Table S5** Pearson regression analysis of global relationships between variables.

**Table S6** Pearson regression analysis of within temperature relationships between stomatal and vein traits with gas exchange.

**Table S7** Pearson regression analysis of within temperature relationships between leaf anatomy and gas exchange.

**Fig. S1** The average environmental conditions recorded in the glasshouse chambers over a day.

**Fig. S2** Representative images of different sorghum lines at the time of measurement.

**Fig. S3** Confocal images to illustrate sampling of stomatal features.

**Fig. S4** Light microscopy image of a cleared and stained leaf.

**Fig. S5** Light microscopy image of a cleared transverse leaf cross-section showing the main anatomical components that were measured.

**Fig. S6** Plots of data points from different sampling times and techniques.

**Fig. S7** Relationships between stomatal anatomy, vein anatomy and leaf width.

**Fig. S8** Relationships between leaf width, interveinal distance and inner leaf anatomy.

**Fig. S9** Structural path analysis of the effect of growth temperature and anatomy on leaf gas exchange.

