## Supplementary for "Anatomical drivers of stomatal conductance in sorghum lines with different leaf widths grown under different temperatures"

Article submission date: 16^th^ October 2022

**Method S1**

The equation for *d* was based on Marrero & Mason (1972):

$ln \left( Pd \right)=\ln\left( A \right)+sln\left( T \right)-\frac{S}{T}$	Eqn. S1

where *A*, *s* and *S* are empirical constants for the diffusion of water vapour in air with *A* = 0.00000187 (atm cm^2^ s^-1^ (K)^-s^); *s* = 2.072; *S* = 0 (K), *T* is the temperature in Kelvin degrees (K) and *P* is atmospheric pressure at our site (102610 Pa). Using eqn. 3 and the value for constants reported before, the units of *d* are in cm^2^ s^-1^, and to be used in eqn. 2, *d* needs to be transformed to m^2^ s^-1^ by dividing by 10000. The calculation for *v* was based on the molar volume of an ideal gas:

$v=\frac{RT}{P}$	Eqn. S2

where *R* is the ideal gas constant = 8.314462618 J mol^-1^ K^-1^. These calculations are explained further in Pan *et al.* (2021).

| **Table S1**. Mean (±SE) of leaf gas exchange, leaf mass per area and leaf width parameters for each *Sorghum* line at the corresponding temperature treatment. The table shows the results of the *post hoc* Tukey’s test for analysis of variance between lines per treatments and between treatments in the last column. Mean values that share similar symbols (lower case letters for within line analysis; upper case letters for within treatment) have no significant difference (*P<*0.05) between them (*n=3*). | | | | | | | | | | | | |
| --- | --- | --- | --- | --- | --- | --- | --- | --- | --- | --- | --- | --- |
| **Unique Line ID** | | FF_SC842-14E | FF_SC449-14E | LR9198 | FF_SC1201-6-3 | FF_SC56-14E | QL12 | R931945-2-2 | SC1079-11Ebk | FF_SC906-14E | FF_SC500-9 | **Treatment Mean** |
| *A_n_ (µmol m^-2^ s^-1^)* | 22°C | 38.9 (1.7) | 39.0 (0.9) | 32.3 (1.7) | 38.5 (3.3) | 32.9 (1.7) | 40.8 (0.6) | 37.8 (1.2) | 36.7 (0.2) | 39.7 (2.9) | 32.6 (1.9) | 36.9 (0.9)A |
|  | 28°C | 38.3 (0.7)a | 47.0 (0.3)bc | 41.9 (2.2)ac | 48.9 (0.3)c | 43.1 (1.4)ac | 48.7 (1.2)c | 43.3 (1.9)ac | 41.1 (0.9)ab | 43.8 (0.4)ac | 42.6 (2.0)ac | 43.9 (1.0)B |
|  | 35°C | 45.8 (0.9)ab | 50.2 (1.1)bc | 43.9 (1.9)a | 51.9 (0.6)c | 51.2 (0.9)bc | 50.5 (0.2)bc | 49.1 (0.6)ac | 45.9 (0.5)ab | 44.2 (0.9)a | 48.3 (1.8)ac | 48.1 (0.9)C |
| *g_s_ (mol m^-2^ s^-1^)* | 22°C | 0.24 (0)ab | 0.23 (0.01)ab | 0.19 (0.01)a | 0.27 (0.03)ab | 0.2 (0.02)a | 0.26 (0.01)ab | 0.24 (0.01)ab | 0.24 (0.01)ab | 0.28 (0.02)b | 0.20 (0.01)a | 0.24 (0.01)A |
|  | 28°C | 0.25 (0.01)a | 0.31 (0)ac | 0.29 (0.03)ac | 0.35 (0.01)c | 0.31 (0.01)ac | 0.31 (0.01)ac | 0.31 (0.01)ac | 0.31 (0.01)ac | 0.33 (0.01)bc | 0.27 (0.01)ab | 0.31 (0.01)B |
|  | 35°C | 0.32 (0.01)a | 0.34 (0.01)ab | 0.33 (0.01)a | 0.43 (0.04)b | 0.39 (0.01)ab | 0.35 (0)ab | 0.39 (0.02)ab | 0.37 (0.01)ab | 0.34 (0.01)a | 0.35 (0.02)ab | 0.36 (0.01)C |
| *iWUE (µmol CO_2_ mol^-1^ H_2_O)* | 22°C | 163.8 (5.3) | 166.9 (1.9) | 169.0 (6.8) | 146.6 (6.7) | 167.4 (9.8) | 159.6 (3.0) | 154.9 (2.3) | 150.9 (3.8) | 139.7 (2.2) | 162.6 (11.2) | 158.2 (2.9)A |
|  | 28°C | 150.8 (2.1) | 150.2 (1.0) | 143.9 (15.9) | 139.9 (2.9) | 137.3 (1.8) | 159.9 (2.5) | 140.9 (5.4) | 132.4 (4.1) | 131.1 (3.0) | 158.1 (1.5) | 144.5 (3.0)B |
|  | 35°C | 142.7 (3.5) | 147.0 (5.7) | 135.0 (5.0) | 125.3 (13.5) | 128.3 (3.4) | 145.9 (1.1) | 124.1 (4.9) | 124.3 (1.8) | 131.1 (3.6) | 136.8 (1.6) | 134.1 (2.7)C |
| *Leaf mass per area (g m^-2^)* | 22°C | 20.6 (0.5)a | 18.6 (2.2) | 23.2 (0.8) | 22.4 (0.6) | 20.9 (0.5) | 25.0 (0.1) | 20.8 (2.1) | 22.3 (0.9) | 23.2 (1.4)a | 22.7 (0.9) | 21.9 (0.5)A |
|  | 28°C | 22.1 (0.7)ab | 29.7 (0.1)e | 21.5 (0.5)a | 27.1 (0.8)de | 22.5 (1.1)ac | 26.9 (0.7)de | 25.8 (0.7)bce | 24.5 (0.3)acd | 26.5 (1.2)ce | 24.5 (1.00acd | 25.1 (0.8)B |
|  | 35°C | 23.6 (0.2)a | 26.9 (0.7)ab | 23.7 (1.2)a | 31.4 (0.8)c | 26.9 (1.1)ab | 30.3 (0)bc | 31.3 (0.4)c | 29.7 (1.2)bc | 27.9 (0.6)bc | 31.4 (0.3)c | 28.3 (0.9)C |
| *Leaf Width (cm)* | 22°C | 2.4 (0.2)a | 2.9 (0.4)ab | 2.7 (0.3)a | 4.6 (0.2)c | 2.8 (0.3)a | 3.4 (0.1)ac | 3.4 (0.4)ac | 4.4 (0.3)bc | 4.7 (0.3)c | 2.6 (0.3)a | 3.4 (0.3)A |
|  | 28°C | 3.4 (0.03)a | 5.5 (0.2)bd | 5.2 (0.4)bc | 6.6 (0.1)d | 5.5 (0.2)bd | 4.5 (0.3)ab | 5.4 (0.2)bd | 5.7 (0.4)bd | 6.2 (0.2)cd | 4.8 (0.2)b | 5.3 (0.3)B |
|  | 35°C | 3.2 (0.1)a | 5.2 (0.2)b | 5.0 (0.2)b | 8.1 (0.1)e | 6.2 (0.2)c | 5.2 (0.02)b | 6.5 (0.1)c | 7.3 (0.2)de | 6.6 (0.3)cd | 6.4 (0.1)c | 5.9 (0.4)C |
| *A_n_*: carbon assimilation rate; *g_s_*: stomatal conductance; *iWUE*: Intrinsic water use efficiency. | | | | | | | | | | | | |

| **Table S2.** Means (±SE) of leaf stomatal parameters for each Sorghum genotype at the corresponding temperature treatment. The table shows the results of the post hoc Tukey’s test for analysis of variance between genotypes per treatment and between treatments in the last column. Mean values that share similar symbols (lower case letters for within genotype analysis; upper case letters for within treatment) have no significant difference (*P*<0.05) between them (*n*=3). Epidermal cell size was measured on seven selected genotypes as this measurement occurred later in the analysis period where time was of constraint. | | | | | | | | | | | | |
| --- | --- | --- | --- | --- | --- | --- | --- | --- | --- | --- | --- | --- |
| **Unique Line ID** | | FF_SC842-14E | FF_SC449-14E | LR9198 | FF_SC1201-6-3 | FF_SC56-14E | QL12 | R931945-2-2 | SC1079-11Ebk | FF_SC906-14E | FF_SC500-9 | **Treatment Mean** |
| *SD (mm^-2^)* | 22°C | 184.1 (9.2)abc | 189.7 (9.9)ad | 149.2 (7.3)a | 220.2 (6.9)cd | 234.4 (11.5)d | 197.5 (10.5)bd | 198.2 (6.1)bd | 151.2 (1.6)a | 169.0 (8.9)ab | 172.8 (5.2)ab | 186.7 (8.3) |
|  | 28°C | 192.9 (9.2) | 199.7 (15.2) | 157.4 (6.6) | 183.8 (3.2) | 209.7 (20.9) | 208.9 (35.4) | 189.9 (20.7) | 174.1 (24.7) | 164.9 (7.5) | 189.9 (17.9) | 187 (5.3) |
|  | 35°C | 191.1 (15.3)ac | 228.3 (5.1)c | 161.1 (6.2)ab | 144.3 (2.6)a | 196.6 (17.7)ac | 209.8 (8.7)bc | 185.8 (8.9)ac | 145.6 (12.3)a | 190.0 (12.1)ac | 233.7 (4.4)c | 188.6 (9.3) |
| *SS (µm^2^)* | 22°C | 741.6 (61)ab | 724.3 (33)a | 1213 (18)d | 1004 (27)cd | 1051 (49)cd | 888.3 (56)ac | 959.0 (15)bc | 1012 (57)cd | 978.1 (14)c | 1045 (48)cd | 961.6 (43.9) |
|  | 28°C | 723.7 (16)a | 1077 (107)b | 1049 (24)b | 996.9 (23)ab | 959.5 (20)ab | 900.7 (54)ab | 1009 (9.4)b | 991.1 (83)ab | 1111 (56)b | 983.4 (19)ab | 980 (32.4) |
|  | 35°C | 695 (23)a | 969 (37)bc | 1002 (51)bc | 1052 (7.6)bc | 1057 (45)bc | 957 (19)bc | 1147 (46)cd | 1265 (23)d | 1041 (52)bc | 922 (33)b | 1010.7 (44.8) |
| *a_max_ (µm^2^)* | 22°C | 142.9 (3.1)a | 144.1 (12.6)a | 214.9 (4.4)a | 215.7 (5.1) | 215.6 (1.6) | 163.9 (15) | 190.2 (4.2) | 188.7 (3.6) | 184.8 (1.3) | 228.7 (15) | 188.9 (9.2)A |
|  | 28°C | 179.2 (6.5)b | 211.1 (18)b | 198.6 (9.5)ab | 222.7 (4.1) | 214.4 (4.2) | 173.2 (12) | 195.4 (9.2) | 206.3 (14) | 202.9 (9.3) | 200.1 (4.6) | 200.4 (4.6)AB |
|  | 35°C | 155.6 (8.7)ab | 195.9 (5.3)ab | 181.1 (7.6)b | 212.4 (3.6) | 224.5 (17) | 181.8 (0.9) | 228.6 (13)a | 245.6 (1.9)a | 205.8 (6.3) | 197.6 (3.5) | 202.9 (7.9)B |
| *Epidermal cell size (µm^2^)* | 22°C | 2830 (665.3) | 2275.8 (191.4) | - | 2130.4 (84.5) | - | 2113 (24.5) | 1912 (22.3) | 3003.4 (253) | 2446.3 (61.2) | - | 2387.3 (139.9) |
|  | 28°C | 2497.9 (31.9)ab | 2593.8 (341)ab | - | 2385.8 (236)ab | - | 2255.2 (237.7)ab | 1831.6 (123.1)a | 2806.4 (364.8)av | 3283.5 (64.7)b | - | 2522 (158.6) |
|  | 35°C | 1848.1 (86.3)a | 2175.2 (214.7)ab | - | 2623.6 (39.2)ab | - | 2366.3 (106.2)ab | 2384.2 (149.5)ab | 2945.8 (262)b | 2742.8 (82.1)ab | - | 2440.8 (128.6) |
| *Operational aperture (µm^2^)* | 22°C | 12.5 (0.2)ab | 12.7 (0.7)ab | 15.7 (1.1)ab | 14.9 (2.4)ab | 10.4 (1.9)a | 14.7 (1.3)ab | 14.7 (0.2)ab | 18.2 (0.5)ab | 19.7 (2.3)b | 14.8 (1.7)ab | 14.8 (0.8)A |
|  | 28°C | 14.7 (1.5) | 19.5 (2.0) | 24.4 (4.6) | 24.5 (0.6) | 19.2 (1.9) | 17.8 (2.8) | 20.5 (1.8) | 23.7 (4.1) | 24.5 (0.7) | 17.6 (1.0) | 20.6 (1.0)B |
|  | 35°C | 19 (1.6)a | 17.8 (0.6)a | 25.3 (2.3)ab | 42.2 (5.8)c | 26.9 (2.9)ab | 18.7 (1.2)a | 27.8 (3.3)ab | 35.3 (3.0)bc | 20.9 (1.8)a | 18.2 (1.2)a | 25.2 (2.5)B |
| *% stomatal aperture* | 22°C | 8.7 (0.2)ac | 8.9 (0.3)ac | 7.3 (0.5)ac | 7.0 (1.2)ac | 4.8 (0.9)a | 9.2 (1.3)bc | 7.7 (0.3)ac | 9.7 (0.1)bc | 10.6 (1.3)c | 6.4 (0.4)ab | 8.03 (0.5)A |
|  | 28°C | 8.2 (0.9) | 9.2 (0.2) | 12.1 (1.7) | 11.0 (0.2) | 9.0 (0.9) | 10.1 (1.3) | 10.5 (1.0) | 11.2 (1.3) | 12.1 (0.3) | 8.8 (0.5) | 10.2 (0.4)AB |
|  | 35°C | 12.1 (0.5) | 9.1 (0.4) | 13.9 (0.7) | 19.8 (2.4)a | 11.9 (0.4) | 10.3 (0.7) | 12.0 (0.8) | 14.4 (1.1) | 10.1 (0.7) | 9.2 (0.5) | 12.3 (0.9)B |
| *g_smax_ (mol m^-2^ s^-1^)* | 22°C | 1.6 (0.1)a | 1.6 (0.04)a | 1.5 (0.1)a | 2.2 (0.1)bc | 2.4 (0.1)c | 1.7 (0.1)a | 1.9 (0.1)ab | 1.5 (0.02)a | 1.6 (0.1)a | 1.8 (0.02)a | 1.8 (0.1) |
|  | 28°C | 1.8 (0.1) | 2.0 (0.1) | 1.6 (0.04) | 1.9 (0.1) | 2.2 (0.2) | 1.9 (0.3) | 1.9 (0.2) | 1.8 (0.2) | 1.7 (0.1) | 1.9 (0.2) | 1.9 (1.0) |
|  | 35°C | 1.7 (0.1)abc | 2.3 (0.04)d | 1.5 (0.03)a | 1.5 (0.02)a | 2.1 (0.1)d | 2.1 (0.1)cd | 2.1 (0.03)bd | 1.7 (0.1)ab | 2.0 (0.1)bd | 2.3 (0.04)d | 1.9 (0.1) |
| *f_gc_ (%)* | 22°C | 6.76 (0.44)a | 6.82 (0.1)ab | 9.04 (0.51)c | 11.03 (0.14)de | 12.23 (0.08)e | 8.77 (0.82)bc | 9.5 (0.36)cd | 7.64 (0.44)ac | 8.28 (0.65)ac | 8.99 (0.18)c | 8.91 (0.55) |
|  | 28°C | 7 (0.56)a | 10.51 (0.35)b | 8.24 (0.28)ab | 9.17 (0.43)ab | 10 (0.97)ab | 9.13 (1.16)ab | 9.57 (1.24)ab | 8.32 (0.5)ab | 9.12 (0.51)ab | 9.29 (0.84)ab | 9.04 (0.31) |
|  | 35°C | 6.59 (0.4)a | 11.05 (0.46)d | 8.03 (0.21)ac | 7.6 (0.22)av | 10.32 (0.95)d | 10.01 (0.26)cd | 10.6 (0.08)d | 9.16 (0.72)bcd | 9.84 (0.71)cd | 10.78 (0.56)d | 9.39 (0.48) |
| *SD*: Stomatal Density; *SS*: Stomatal Size; *a_max_*: maximum stomatal aperture; *g_smax_*: theoritical anatomical conductance; *f_gc_*: fraction of epidermis allocated to stomata. | | | | | | | | | | | | |

| **Table S3**. Means (±SE) of leaf vein anatomical parameters for each *Sorghum* line at the corresponding temperature treatment. The table shows the results of the post hoc Tukey’s test for analysis of variance between lines per treatments and between treatments in the last column. Mean values that share similar symbols (lower case letters for within line analysis; upper case letters for within treatment) have no significant difference (*P<*0.05) between them (*n=3*). | | | | | | | | | | | | | | | | | | | | | | |
| --- | --- | --- | --- | --- | --- | --- | --- | --- | --- | --- | --- | --- | --- | --- | --- | --- | --- | --- | --- | --- | --- | --- |
| **Unique Line ID** | | | FF_SC842-14E | FF_SC449-14E | | LR9198 | | FF_SC1201-6-3 | | FF_SC56-14E | | QL12 | | R931945-2-2 | | SC1079-11Ebk | | FF_SC906-14E | | FF_SC500-9 | | **Treatment Mean** |
| *Interveinal Distance (µm)* | 22°C | | 127.1 (2.8)ab | 116.7 (2.7)a | | 127.2 (2.8)ab | | 117.4 (9.7)ab | | 124.1 (5.4)ab | | 133.9 (8.3)ab | | 127.3 (3.9)ab | | 134.0 (4.6)ab | | 149.9 (6.6)b | | 134.2 (9.7)ab | | 129.2 (2.3)A |
|  | 28°C | | 131.2 (1.3) | 119.4 (3.1) | | 120.1 (6.3) | | 110.8 (8.8) | | 102.7 (0.1) | | 120.6 (7.8) | | 114.1 (3.2) | | 119.4 (5.8) | | 126.8 (10.2) | | 124.2 (3.7) | | 119 (2.5)B |
|  | 35°C | | 116.6 (4.6)ab | 109.5 (5.7)a | | 114.4 (2.9)a | | 127.8 (5.9)ab | | 123.7 (4.4)ab | | 120.4 (6.9)ab | | 144.9 (5.9)b | | 129.1 (7.7)ab | | 124.5 (4.5)ab | | 124.9 (3.8)ab | | 123.6 (2.9)AB |
| *VD (mm mm^-2^)* | 22°C | | 10.5 (0.1) | 11.1 (0.3) | | 10.8 (0.1) | | 11.8 (0.7) | | 11.5 (0.4) | | 11.3 (0.5) | | 11.2 (0.3) | | 10.4 (0.2) | | 9.8 (0.4) | | 10.2 (0.5) | | 10.9 (0.2)A |
|  | 28°C | | 10.5 (0.3) | 11.6 (0.2) | | 11.3 (0.6) | | 11.7 (0.7) | | 12.7 (0.02) | | 11.7 (0.5) | | 11.6 (0.2) | | 11.1 (0.5) | | 11.3 (0.6)a | | 11.6 (0.3) | | 11.5 (0.2)B |
|  | 35°C | | 11.5 (0.4)ab | 12.5 (0.4)b | | 11.8 (0.1)ab | | 10.6 (0.60)ab | | 11.1 (0.4)ab | | 11.9 (0.6)ab | | 10.1 (0.3)a | | 10.9 (0.3)ab | | 11.5 (0.4)ab | | 11.6 (0.3)ab | | 11.3 (0.2)AB |
| *Total Number of Longitudinal Veins* | 22°C | | 156.4 (1.8)a | 195.8 (25.8)ab | | 423.3 (16.6)e | | 423.7 (9.3)e | | 369.5 (13.9)de | | 319.8 (24.1)cd | | 326 (11.5)cd | | 291.6 (15.3)cd | | 254.9 (28.7)bc | | 334.2 (0)ce | | 309.5 (26.5)A |
|  | 28°C | | 247.1 (21.5)a | 419.5 (27.9)b | | 432.6 (33)b | | 416.8 (16.4)b | | 375.3 (8.8)ab | | 364.8 (14.6)ab | | 311.1 (6.6)ab | | 419 (42.6)b | | 422.7 (0)b | | 393.4 (0)ab | | 380.2 (17.8)B |
|  | 35°C | | 230.3 (3.7)a | 428.8 (17.2)bcd | | 478.9 (16.5)cd | | - | | 433.7 (4.6)bcd | | 357.5 (7.5)ac | | 437.7 (20.6)bcd | | 503.3 (36.1)d | | 449.1 (29.1)cd | | 304.8 (3.3)ab | | 402.7 (27.7)B |
| *VD*: Vein density. | | | | | | | | | | | | | | | | | | | | | | |

| **Table S4.** Means (±SE) of measured inner leaf anatomical parameters for each Sorghum genotype at the corresponding temperature treatment. The table shows the results of the post hoc Tukey’s test for analysis of variance between genotypes per treatments and between treatments in the last column. Mean values that share similar symbols ( lower case letters for within genotype analysis; upper case letters for within treatment) have no significant difference (*P*<0.05) between them (*n*=3). | | | | | | | | | | | | |
| --- | --- | --- | --- | --- | --- | --- | --- | --- | --- | --- | --- | --- |
| **Unique Genotype ID** | | FF_SC842-14E | FF_SC449-14E | LR9198 | FF_SC1201-6-3 | FF_SC56-14E | QL12 | R931945-2-2 | SC1079-11Ebk | FF_SC906-14E | FF_SC500-9 | **Treatment Mean** |
| *MT (µm)* | 22°C | 100.4 (2.0)ab | 97.1 (2.6)a | 122.0 (8.6)bc | 110.6 (0)ac | 97.3 (8.4)ab | 97.5 (5.7)abd | 118.8 (5.5)ac | 129.1 (5.5)c | 131.6 (1.2)c | 131.1 (4.1)dc | 113.6 (4.4)AB |
|  | 28°C | 91.3 (2.5)abc | 126.4 (4.3)cd | 118.1 (5.5)ad | 94.9 (1.2)abc | 82.3 (7.9)a | 118.1 (0.7)ad | 88.9 (6.8)ab | 118.9 (9.1)bd | 132.5 (6.8)d | 103.9 (7.3)ad | 107.5 (5.2)A |
|  | 35°C | 91.3 (0)a | 107.0 (3.3)a | - | 115.9 (0.3)a | 124.4 (7.9)a | 112.9 (4.1)a | 136.4 (2.5)ab | 160.7 (3.6)b | 127.3 (4.3)a | 108.2 (8.5)a | 120.5 (8.3)B |
| *M / BS ratio* | 22°C | 2.5 (0.2)ab | 2.9 (0.3)b | 2.5 (0.1)ab | 2.5 (0.1)ab | 1.9 (0.1)a | 1.9 (0.2)a | 2.2 (0.2)ab | 2.4 (0.1)ab | 2.1 (0)a | 2.1 (0.1)ab | 2.3 (0.1) |
|  | 28°C | 3.1 (0.1)d | 3.5 (0.1)d | 2.9 (0.1)cd | 2.2 (0.2)ac | 1.7 (0.2)ab | 2.3 (0.1)bc | 1.6 (0.1)a | 2.2 (0.1)ac | 2.1 (0.1)ab | 2.1 (0.1)ac | 2.4 (0.2) |
|  | 35°C | 2.9 (0)ab | 3.5 (0.1)b | - | 2.9 (0.7)ab | 2.5 (0.1)ab | 2.5 (0.1)ab | 2.3 (0.1)a | 2.3 (0.1)a | 2.4 (0.1)a | 1.8 (0.1)a | 2.6 (0.2) |
| *M_si_ (µm^2^ µm^-1^)* | 22°C | 53.5 (2.2)bc | 55.2 (2.9)ac | 61.8 (3.2)c | 55.4 (1.3)bc | 43.2 (4.4)ab | 38.0 (0.6)b | 52.9 (3.1)bc | 61.7 (3.4)c | 58.4 (0.4)c | 53.4 (0.4)bc | 53.4 (2.3) |
|  | 28°C | 50.4 (0.8)bc | 74.5 (0.3)d | 61.4 (4.5)cd | 45.8 (0.5)ac | 32.8 (4.3)a | 52.8 (1.7)bc | 36.7 (5.2)ab | 54.4 (4.6)c | 55.9 (0.5)cd | 42.1 (0.9)ac | 50.7 (3.7) |
|  | 35°C | 53.1 (0)ab | 64.9 (2.9)b | - | 50.8 (7.0)ab | 52.7 (0.8)ab | 50.9 (2.7)ab | 59.7 (0.9)b | 67.2 (1.2)b | 55.6 (2.1)ab | 38.8 (3.9)a | 54.9 (2.7) |
| *BS_si_ (µm^2^ µm^-1^)* | 22°C | 21.7 (1.1)ab | 19.2 (1.3)a | 24.8 (0.7)ac | 22.1 (0.5)ac | 21.7 (1.8)ab | 20.2 (1.7)ab | 24.4 (2.2)ac | 26.5 (0.2)bc | 28.4 (0.7)c | 25.3 (0.8)ac | 23.4 (0.9) |
|  | 28°C | 16.5 (0.4)a | 21.5 (0.6)ac | 21.4 (1.0)ac | 21.5 (1.1)ab | 19.0 (0.5)ab | 22.8 (0.1)bc | 22.8 (1.2)bc | 24.4 (1.1)abc | 26.9 (1.2)c | 20.2 (1.7)ab | 21.7 (0.9) |
|  | 35°C | 18.6 (0)ab | 18.8 (0.4)b | - | 18.9 (1.0)b | 21.7 (1.2)ab | 20.7 (0.6)ab | 25.8 (0.9)ac | 29.9 (1.0)c | 23.4 (0.7)ab | 22.1 (1.3)ab | 22.2 (1.2) |
| *VB_si_ (µm^2^ µm^-1^)* | 22°C | 4.7 (0.4)ab | 4.3 (0.6)a | 7.2 (0.3)b | 5.5 (0.7)ab | 5.7 (0.6)ab | 5.3 (0.3)ab | 6.9 (0.7)b | 5.4 (0.4)ab | 5.9 (0.6)ab | 6.1 (0.6)ab | 5.7 (0.3) |
|  | 28°C | 4.7 (0.5) | 6.8 (0.3)b | 6.8 (1.0) | 4.6 (0.2) | 4.9 (0.5) | 5.5 (0.2) | 5.7 (0.5)a | 5.1 (0.7) | 5.8 (0.3) | 6.3 (0.4) | 5.6 (0.3) |
|  | 35°C | 3.9 (0) | 5.4 (0.3) | - | 4.9 (0.1) | 6.3 (0.6) | 4.7 (0.4) | 7.6 (0.5) | 7.4 (0.3) | 6.0 (0.4) | 7.8 (1.4) | 6.0 (0.4) |
| *IAS_si_ (µm^2^ µm^-1^)* | 22°C | 17.3 (0.6)ab | 13.8 (0.4)a | 27.3 (8.3)ac | 23.9 (0.7)ac | 24.1 (1.2)ac | 31.3 (4.2)bc | 26.8 (1.0)ac | 29.1 (1.7)ac | 31.4 (0.4)bc | 42.6 (4.8)c | 26.8 (2.4)A |
|  | 28°C | 14.9 (1.0)a | 16.8 (2.5)a | 26.5 (0.7)ab | 19.1 (0.5)a | 23.9 (4.5)ab | 34.8 (3.5)ab | 21.9 (1.7)ab | 30.7 (2.9)ab | 40.4 (7.9)b | 29.1 (4.5)ab | 25.8 (2.4)A |
|  | 35°C | 11.9 (0)a | 17.8 (1.4)a | - | 33.6 (2.9)ab | 39.1 (7.8)ab | 31.9 (1.4)ab | 34.4 (2.0)ab | 51.5 (2.6)b | 38.1 (1.9)ab | 35.3 (2.7)ab | 32.6 (3.7)B |
| *M cell length (µm)* | 22°C | 21.8 (0.8)ab | 21.7 (0.8)ab | 20.8 (1.6)ab | 20.7 (1.2)ab | 17.4 (1.6)a | 17.6 (0.1)a | 21.6 (0.4)ab | 24.2 (1.6)b | 24.4 (0.4)b | 21.2 (0.6)ab | 21.1 (0.7)A |
|  | 28°C | 19.5 (0.7)bcd | 23.9 (0.8)d | 21 (0.4)bcd | 17.4 (0.3)ac | 13.6 (1.1)a | 20.8 (0)bcd | 16.2 (1.2)ab | 20.9 (1.8)bcd | 21.4 (0.7)cd | 18.4 (0.2)ad | 19.3 (0.9)B |
|  | 35°C | 19.4 (0)ab | 21.0 (0.7)ab | - | 19.6 (0.9)ab | 20.3 (0.4)ab | 20.7 (0.6)ab | 25.1 (0.5)cd | 26.3 (0.8)d | 22.1 (0.5)bc | 17.3 (0.7)a | 21.3 (0.9)A |
| *M cell area (µm^2^)* | 22°C | 330.2 (9.3)ab | 283.1 (16.4)ab | 323.3 (19.3)ab | 334.6 (26.9)ab | 208.9 (41.4)a | 195.1 (4.3)a | 257.4 (14.8)ab | 380.7 (37.0)b | 347.8 (6.7)b | 311.6 (24.5)ab | 297.3 (18.1) |
|  | 28°C | 262.9 (1.7)bc | 339.6 (26.2)b | 364.6 (0)b | 229.8 (8.2)ab | 146.7 (20)a | 279.3 (5.8)bc | 169.1 (30)ac | 294.5 (38.2)bc | 318.5 (14.9)b | 231.6 (13.4)ab | 262.8 (17.9) |
|  | 35°C | 241.7 (0)ab | 280.9 (12.1)ab | - | 290.2 (45.4)ab | 267.9 (12.8)ab | 268.6 (13.3)ab | 335.9 (3.5)bc | 413.6 (3.5)c | 310.2 (12.9)ac | 215.5 (23.5)a | 291.6 (18.2) |
| *M*: Mesophyll; *BS*: Bundle Sheath; *MT:* Leaf Mesophyll Thickness; *M_si_*: Mesophyll surface area per interveinal distance; *BS_si_*: Bundle sheath surface area per interveinal distance; *VB_si_*: Vascular bundle surface area per interveinal distance; *IAS_si_*: Intercellular airspace surface area per interveinal distance; *Dist_s-v_*: distance between stomata and nearest projected VB. | | | | | | | | | | | | |

**Table 4 continued…**

| **Unique Genotype ID** | | FF_SC842-14E | FF_SC449-14E | LR9198 | FF_SC1201-6-3 | FF_SC56-14E | QL12 | R931945-2-2 | SC1079-11Ebk | FF_SC906-14E | FF_SC500-9 | **Treatment Mean** |
| --- | --- | --- | --- | --- | --- | --- | --- | --- | --- | --- | --- | --- |
| *BS cell area (µm^2^)* | 22°C | 365.5 (36.1)ab | 301.9 (38.1)a | 377.3 (13.2)ab | 346.7 (17.7)ab | 284.7 (42.8)a | 312.6 (28.6)a | 375.3 (50.6)ab | 485.2 (20.5)bc | 603.1 (12.3)c | 428.4 (10.2)ab | 388.1 (28.9) |
|  | 28°C | 248.6 (10.7)a | 338.7 (21.1)ab | 313.8 (35.5)a | 321.2 (20.0)a | 225.9 (6.5)a | 380.8 (5.2)ab | 299.9 (23.9)a | 382.2 (50.8)ab | 492.3 (32.2)b | 361.1 (50.4)ab | 336.5 (22.6) |
|  | 35°C | 294.5 (0)ac | 265.3 (6.8)a | - | 291.9 (4.6)ab | 312.6 (28.6)ab | 307.7 (2.3)ab | 444.8 (17.3)cd | 530.3 (20.9)d | 398.8 (22.4)bc | 383.3 (33.0)ac | 358.8 (27.5) |
| *Interveinal Distance (µm)* | 22°C | 135.4 (5.4)ab | 126.4 (6.7)a | 132.4 (6.7)a | 139.3 (4.1)ab | 116.2 (6.5)a | 126.1 (0.6)a | 129.6 (8.3)a | 141 (6.4)ab | 157.9 (1.7)b | 137.6 (2.7)ab | 134.2 (3.4)A |
|  | 28°C | 125.2 (3.2)ab | 128.5 (11.3)ab | 132 (8.5)ab | 123.8 (3.1)ab | 102.5 (2.6)a | 131.3 (1.9)ab | 116.2 (4.5)ab | 117.2 (10.7)ab | 142.5 (3.7)b | 143.3 (6.9)b | 126.2 (3.7)B |
|  | 35°C | 123.8 (0)ab | 114.8 (2.6)a | - | 135.9 (3.5)ab | 127.4 (8.1)ab | 116.9 (3.7)a | 145.6 (2.1)b | 143.8 (2.7)b | 136.2 (2.9)ab | 138.8 (4.1)ab | 131.5 (3.5)C |
| *Dist_s-v_ (µm)* | 22°C | 74.87 (3.29) | 73.02 (2.27) | 80.18 (2.67) | 73.06 (0.53) | 72.49 (3.55) | 71.16 (5.14) | 81.03 (2.39) | 85.71 (4.02) | 83.36 (3.09) | 83.28 (1.74) | 77.82 (1.72)A |
|  | 28°C | 71.98 (1.1)a | 81.18 (3.08)ab | 74.85 (0.49)a | 69.38 (3.43)a | 64.1 (3.38)a | 76.44 (0.67)a | 67.19 (3.25)a | 80.31 (5.13)ab | 94.9 (4.92)b | 68.79 (4.16)a | 74.91 (2.84)A |
|  | 35°C | 71.99 (0)b | 74.55 (3.71)b | - | 86.56 (4.13)ab | 80.96 (5.07)b | 76.06 (3.51)b | 89.53 (2.61)ab | 99 (0.93)a | 86.56 (2.88)ab | 79.22 (3.47)b | 82.72 (2.69)B |
| *Hydraulic Distance (µm)* | 22°C | 90.15 (1.42)ab | 87.7 (1.92)a | 100.82 (3.7)ab | 91.65 (0.26)ab | 87.41 (1.13)a | 86.26 (3.65)ac | 100.51 (1.59)ab | 107.3 (0.88)b | 106.22 (0.51)bc | 105.98 (1.92)ab | 96.4 (2.72)A |
|  | 28°C | 85.24 (0.55)a | 102.89 (1.78)ab | 95.39 (1.32)ab | 84.11 (0.7)a | 77.77 (2.93)a | 96.59 (2.42)ab | 80.63 (2.56)a | 99.94 (0.65)ab | 115.75 (2.92)b | 86.2 (2.26)a | 92.5 (3.73)A |
|  | 35°C | 85.26 (0)a | 91.83 (0.39)a | - | 104.23 (6.16)ab | 102.15 (3.78)a | 94.78 (1.54)a | 112.56 (0.94)ab | 127.54 (0.23)b | 107.46 (1.64)ab | 95.99 (2.45)a | 102.4 (3.98)B |
| *M*: Mesophyll; *BS*: Bundle Sheath; *MT:* Leaf Mesophyll Thickness; *M_si_*: Mesophyll surface area per interveinal distance; *BS_si_*: Bundle sheath surface area per interveinal distance; *VB_si_*: Vascular bundle surface area per interveinal distance; *IAS_si_*: Intercellular airspace surface area per interveinal distance; *Dist_s-v_*: distance between stomata and nearest projected VB. | | | | | | | | | | | | |

| **Table S5**. Pearson product-moment correlation analysis results for the relationships between the measured variables. The *R*coefficient and the statistical significance were determined using the mean value per line, per treatment for each variable. Statistical significance was judged as: *P*<0.001 (***), *P*<0.05 (**), *P*<0.1 (*), *P*>0.1 (ns). Underlined coefficients show the correlation between the variables was significant after excluding the 22°C treatment. (*n*=3). | | | | | | | | | | | | | | | | | | | | | | | | | |
| --- | --- | --- | --- | --- | --- | --- | --- | --- | --- | --- | --- | --- | --- | --- | --- | --- | --- | --- | --- | --- | --- | --- | --- | --- | --- |
|  | *IVD_c_* | *M / BS* | *MC_length_* | *MC_area_* | *BSC_area_* | *M_si_* | *BS_si_* | *VB_si_* | *IAS_si_* | *Dist_H_* | *SD* | *SS* | *a_max_* | *ES* | *a_op_* | *% a* | *g_smax_* | *IVD* | *VD* | *TLV* | *LMA* | *A_n_* | *g_s_* | *iWUE* | *LW* |
| *MT* | 0.66*** | ns | 0.83*** | 0.8*** | 0.79*** | 0.68*** | 0.8*** | 0.62*** | 0.75*** | 0.97*** | -0.51** | 0.69*** | 0.49** | 0.59** | 0.46** | 0.39* | ns | 0.55** | -0.48** | 0.49** | 0.37** | ns | 0.46** | -0.23** | 0.5** |
| *IVD_c_* | - | ns | 0.64*** | 0.64*** | 0.82*** | 0.39** | 0.61*** | 0.46** | 0.47** | 0.65*** | -0.45** | 0.52** | ns | 0.44** | ns | ns | -0.36* | 0.48*** | -0.71*** | ns | ns | ns | ns | ns | ns |
| *M / BS* | - | - | 0.39** | ns | ns | 0.68*** | -0.4** | ns | ns | ns | ns | ns | ns | ns | ns | ns | ns | ns | ns | ns | ns | ns | ns | ns | ns |
| *MC_length_* | - | - | - | 0.93*** | 0.7*** | 0.87*** | 0.62*** | 0.34* | ns | 0.8*** | -0.45** | 0.51** | ns | 0.51** | ns | ns | -0.36* | 0.59** | -0.6** | 0.54** | ns | ns | ns | ns | ns |
| *MC_area_* | - | - | - | - | 0.7*** | 0.86*** | 0.67*** | ns | ns | 0.77*** | -0.59** | 0.41** | ns | 0.68*** | 0.48* | ns | -0.44** | 0.47** | -0.57** | 0.56** | ns | ns | ns | ns | ns |
| *BSC_area_* | - | - | - | - | - | 0.41** | 0.91*** | 0.44** | 0.6*** | 0.77*** | -0.48** | 0.43** | ns | 0.53** | ns | ns | -0.34* | 0.69** | -0.59*** | 0.53** | ns | ns | ns | ns | ns |
| *M_si_* | - | - | - | - | - | - | 0.39** | ns | ns | 0.65*** | -0.39** | 0.33* | ns | 0.43* | ns | ns | ns | ns | ns | 0.49** | ns | ns | ns | ns | ns |
| *BS_si_* | - | - | - | - | - | - | - | 0.55** | 0.66*** | 0.78*** | -0.47** | 0.64*** | 0.41** | 0.46** | 0.4* | ns | ns | 0.54** | -0.48** | 0.64** | ns | ns | ns | -0.51** | 0.51** |
| *VB_si_* | - | - | - | - | - | - | - | - | 0.54** | 0.56** | ns | 0.69*** | 0.65** | ns | ns | ns | ns | 0.38** | -0.46* | 0.52** | 0.35* | ns | ns | ns | 0.43* |
| *IAS_si_* | - | - | - | - | - | - | - | - | - | 0.74*** | ns | 0.65*** | 0.55** | 0.49** | 0.49** | 0.39* | ns | 0.43** | -0.41** | 0.51** | 0.46** | ns | 0.46** | -0.43** | 0.59*** |
| *Dist_H_* | - | - | - | - | - | - | - | - | - | - | -0.53** | 0.65** | 0.46** | 0.67*** | 0.52** | 0.38** | ns | 0.57** | -0.5** | 0.44** | 0.41** | ns | 0.52 | -0.44** | 0.41** |
| *SD* | - | - | - | - | - | - | - | - | - | - | - | -0.41** | ns | -0.63** | -0.55** | -0.56** | 0.88*** | -0.35* | 0.57** | -0.47** | ns | ns | ns | 0.48** | ns |
| *SS* | - | - | - | - | - | - | - | - | - | - | - | - | 0.85*** | 0.56** | 0.5** | ns | ns | 0.34* | -0.63** | 0.57** | 0.43** | ns | 0.57** | -0.44** | 0.65*** |
| *a_max_* | - | - | - | - | - | - | - | - | - | - | - | - | - | ns | 0.48** | ns | -0.34* | ns | -0.47** | 0.73*** | 0.42** | ns | 0.64*** | -0.48** | 0.5** |
| *ES* | - | - | - | - | - | - | - | - | - | - | - | - | - | - | ns | ns | -0.38* | ns | ns | ns | ns | ns | ns | -0.48* | 0.38* |
| *a_op_* | - | - | - | - | - | - | - | - | - | - | - | - | - | - | - | 0.93*** | ns | ns | -0.68** | 0.64*** | 0.63*** | 0.62*** | 0.81*** | -0.82*** | 0.84*** |
| *% a* | - | - | - | - | - | - | - | - | - | - | - | - | - | - | - | - | -0.42* | ns | -0.59** | 0.34* | 0.53** | 0.65*** | 0.78*** | -0.74*** | 0.71*** |
| *g_smax_* | - | - | - | - | - | - | - | - | - | - | - | - | - | - | - | - | - | ns | 0.52** | ns | ns | ns | 0.49** | -0.36* | 0.49** |
| *IVD* | - | - | - | - | - | - | - | - | - | - | - | - | - | - | - | - | - | - | -0.87*** | ns | 0.59** | ns | 0.48** | 0.42* | 0.45* |
| *VD* | - | - | - | - | - | - | - | - | - | - | - | - | - | - | - | - | - | - | - | ns | 0.44* | ns | -0.54** | 0.54** | -0.57** |
| *TLV* | - | - | - | - | - | - | - | - | - | - | - | - | - | - | - | - | - | - | - | - | 0.46** | 0.32* | 0.47** | -0.49** | 0.65*** |
| *LMA* | - | - | - | - | - | - | - | - | - | - | - | - | - | - | - | - | - | - | - | - | - | 0.79*** | 0.8*** | -0.62*** | 0.75*** |
| *A_n_* | - | - | - | - | - | - | - | - | - | - | - | - | - | - | - | - | - | - | - | - | - | - | 0.92*** | -0.65*** | 0.72*** |
| *g_s_* | - | - | - | - | - | - | - | - | - | - | - | - | - | - | - | - | - | - | - | - | - | - | - | -0.88*** | 0.87*** |
| *iWUE* | - | - | - | - | - | - | - | - | - | - | - | - | - | - | - | - | - | - | - | - | - | - | - | - | -0.87*** |
| *MT*: leaf mesophyll thickness (μm); *IVD_c_*: interveinal Distance measured from leaf cross-sections (μm); *M : BS*: mesophyll to bundle sheath ratio; *MC_length_*: length of mesophyll cell (μm); *MC_area_*: area of single mesophyll cell (μm^2^); *BSC_area_*: area of single bundle sheath cell (μm^2^); *M_si_*: mesophyll surface area per *IVD_c_* (μm^2^ μm^-1^); *BS_si_*: bundle sheath surface area per *IVD_c_* (μm^2^ μm^-1^); *VB_si_*: vascular bundle surface area per *IVD_c_* (μm^2^ μm^-1^); *IAS_si_*: intercellular airspace surface area per *IVD_c_* (μm^2^ μm^-1^); *Dist_H_*: Hydraulic Distance (μm); *SD*: stomatal density (mm^-2^); *SS*: stomatal size (μm^2^); *a_max_*: maximum stomatal pore aperture (μm^2^); *ES*: Epidermal cell size (μm^2^); *a_op_*: operation stomatal aperture (μm^2^); *% a*: percentage of *a_op_* to *a_max_*; *g_max_*: maximum theoritical stomatal conductance (mol m^-2^ s^-1^); *IVD*: inter-veinal distance (mm); *VD*: total vein density (mm mm^-2^); *TLV*: Total Number of longitudinal veins; *LMA*: leaf mass per area (g m^-2^); *A_n_*: net carbon assimilation rate (μmol m^-2^ s^-1^); *g_s_*: stomatal conductance (mol m^-2^ s^-1^); i*WUE*: intrinsic water use efficiency (µmol CO_2_ mol^-1^ H_2_O); *LW*: leaf width (cm). | | | | | | | | | | | | | | | | | | | | | | | | | |

| **Table S6**. Pearson product-moment correlation analysis results for the relationships between the measured variables at different temperatures. The *R* coefficient and the statistical significance were determined using the mean value per line, per treatment for each variable. Statistical significance was judged as: *P*<0.001 (***), *P*<0.05 (**), *P*<0.1 (*), *P*>0.1 (ns). This table shows the relationships between stomatal, vein and gas exchange parameters. (*n*=3). | | | | | | | | | | | | | | | | | | | | | | | | | | | | | | | |
| --- | --- | --- | --- | --- | --- | --- | --- | --- | --- | --- | --- | --- | --- | --- | --- | --- | --- | --- | --- | --- | --- | --- | --- | --- | --- | --- | --- | --- | --- | --- | --- |
|  | | *ES* | | | *a_op_* | | | *% a* | | | *g_smax_* | | | *IVD* | | | *VD* | | | *A_n_* | | | *g_s_* | | | *iWUE* | | | *LW* | | |
|  |  | 22 | 28 | 35 | 22 | 28 | 35 | 22 | 28 | 35 | 22 | 28 | 35 | 22 | 28 | 35 | 22 | 28 | 35 | 22 | 28 | 35 | 22 | 28 | 35 | 22 | 28 | 35 | 22 | 28 | 35 |
| *SD* | 22 | -0.78** | - | - | -0.67** | - | - | ns | - | - | 0.88*** | - | - | 0.77** | - | - | 0.76** | - | - | ns | - | - | ns | - | - | ns | - | - | ns | - | - |
|  | 28 | - | ns | - | - | -0.73** | - | - | -0.78** | - | - | 0.88*** | - | - | 0.7** | - | - | ns | - | - | ns | - | - | ns | - | - | ns | - | - | ns | - |
|  | 35 | - | - | ns | - | - | -0.85** | - | - | -0.87*** | - | - | 0.91*** | - | - | ns | - | - | 0.57* | - | - | ns | - | - | ns | - | - | 0.67** | - | - | ns |
| *SS* | 22 | ns | - | - | ns | - | - | ns | - | - | ns | - | - | ns | - | - | ns | - | - | -0.83** | - | - | ns | - | - | ns | - | - | ns | - | - |
|  | 28 | - | ns | - | - | 0.75** | - | - | 0.62* | - | - | ns | - | - | ns | - | - | ns | - | - | ns | - | - | 0.66** | - | - | ns | - | - | 0.8** | - |
|  | 35 | - | - | 0.87** | - | - | 0.65* | - | - | - | - | - | ns | - | - | 0.59* | - | - | -0.67** | - | - | ns | - | - | 0.55* | - | - | -0.76** | - | - | 0.78** |
| *a_max_* | 22 | ns | - | - | ns | - | - | 6 | - | - | ns | - | - | ns | - | - | ns | - | - | -0.75** | - | - | ns | - | - | ns | - | - | ns | - | - |
|  | 28 | - | ns | - | - | ns | - | - | ns | - | - | ns | - | - | ns | - | - | ns | - | - | ns | - | - | 0.64* | - | - | -0.57* | - | - | 0.79** | - |
|  | 35 | - | - | ns | - | - | 0.59* | - | - | ns | - | - | ns | - | - | 0.66* | - | - | -0.62* | - | - | ns | - | - | 0.7** | - | - | -0.79** | - | - | 0.69** |
| *ES* | 22 | - | - | - | ns | - | - | ns | - | - | ns | - | - | ns | - | - | ns | - | - | ns | - | - | ns | - | - | ns | - | - | ns | - | - |
|  | 28 | - | - | - | - | ns | - | - | ns | - | - | ns | - | - | 0.74* | - | - | ns | - | - | ns | - | - | ns | - | - | ns | - | - | ns | - |
|  | 35 | - | - | - | - | - | ns | - | - | ns | - | - | ns | - | - | ns | - | - | ns | - | - | ns | - | - | ns | - | - | -0.7* | - | - | 0.88** |
| *a_op_* | 22 | - | - | - | - | - | - | 0.67** | - | - | ns | - | - | 0.72** | - | - | -0.71** | - | - | ns | - | - | ns | - | - | -0.75** | - | - | 0.73** | - | - |
|  | 28 | - | - | - | - | - | - | - | 0.91*** | - | - | ns | - | - | ns | - | - | ns | - | - | ns | - | - | 0.77** | - | - | -0.72** | - | - | 0.85** | - |
|  | 35 | - | - | - | - | - | - | - | - | 0.92*** | - | - | -0.65** | - | - | ns | - | - | -0.72** | - | - | ns | - | - | 0.77** | - | - | -0.79** | - | - | 0.73*** |
| *% a* | 22 | - | - | - | - | - | - | - | - | - | -0.74** | - | - | ns | - | - | ns | - | - | 0.73** | - | - | 0.71** | - | - | ns | - | - | ns | - | - |
|  | 28 | - | - | - | - | - | - | - | - | - | - | -0.65** | - | - | ns | - | - | ns | - | - | ns | - | - | 0.62* | - | - | -0.58* | - | - | 0.63* | - |
|  | 35 | - | - | - | - | - | - | - | - | - | - | - | -0.83** | - | - | ns | - | - | -0.57* | - | - | ns | - | - | 0.59* | - | - | -.0.58* | - | - | ns |
| *SD*: stomatal density (mm^-2^); *SS*: stomatal size (μm^2^); *a_max_*: maximum stomatal pore aperture (μm^2^); *ES*: Epidermal cell size (μm^2^); *a_op_*: operation stomatal aperture (μm^2^); *% a*: percentage of *a_op_* to *a_max_*; *g_max_*: maximum theoritical stomatal conductance (mol m^-2^ s^-1^); *IVD*: inter-veinal distance (mm); *VD*: total vein density (mm mm^-2^); *A_n_*: net carbon assimilation rate (μmol m^-2^ s^-1^); *g_s_*: stomatal conductance (mol m^-2^ s^-1^); i*WUE*: intrinsic water use efficiency (µmol CO_2_ mol^-1^ H_2_O); *LW*: leaf width (cm). | | | | | | | | | | | | | | | | | | | | | | | | | | | | | | | |

**Table 6 continued…**

|  | | *IVD* | | | *VD* | | | *A_n_* | | | *g_s_* | | | *iWUE* | | | *LW* | | |
| --- | --- | --- | --- | --- | --- | --- | --- | --- | --- | --- | --- | --- | --- | --- | --- | --- | --- | --- | --- |
|  |  | 22 | 28 | 35 | 22 | 28 | 35 | 22 | 28 | 35 | 22 | 28 | 35 | 22 | 28 | 35 | 22 | 28 | 35 |
| *g_smax_* | 22 | ns | - | - | 0.64* | - | - | ns | - | - | ns | - | - | ns | - | - | ns | - | - |
|  | 28 | - | -0.67* | - | - | 0.8** | - | - | ns | - | - | ns | - | - | ns | - | - | ns | - |
|  | 35 | - | - | ns | - | - | ns | - | - | ns | - | - | ns | - | - | ns | - | - | ns |
| *IVD* | 22 | - | - | - | -0.8** | - | - | ns | - | - | ns | - | - | ns | - | - | ns | - | - |
|  | 28 | - | - | - | - | -0.79** | - | - | ns | - | - | ns | - | - | ns | - | - | ns | - |
|  | 35 | - | - | - | - | - | -0.92*** | - | - | ns | - | - | 0.62* | - | - | -0.75** | - | - | 0.6* |
| *VD* | 22 | - | - | - | - | - | - | ns | - | - | ns | - | - | ns | - | - | ns | - | - |
|  | 28 | - | - | - | - | - | - | - | ns | - | - | ns | - | - | ns | - | - | ns | - |
|  | 35 | - | - | - | - | - | - | - | - | ns | - | - | -0.78** | - | - | 0.9*** | - | - | -0.72** |
| *An* | 22 | - | - | - | - | - | - | - | - | - | 0.88*** | - | - | ns | - | - | ns | - | - |
|  | 28 | - | - | - | - | - | - | - | - | - | - | 0.66** | - | - | ns | - | - | ns | - |
|  | 35 | - | - | - | - | - | - | - | - | - | - | - | 0.69** | - | - | ns | - | - | ns |
| *gs* | 22 | - | - | - | - | - | - | - | - | - | - | - | - | -0.85** | - | - | 0.78** | - | - |
|  | 28 | - | - | - | - | - | - | - | - | - | - | - | - | - | -0.6* | - | - | 0.91*** | - |
|  | 35 | - | - | - | - | - | - | - | - | - | - | - | - | - | - | -0.71** | - | - | 0.75** |
| *iWUE* | 22 | - | - | - | - | - | - | - | - | - | - | - | - | - | - | - | -0.92*** | - | - |
|  | 28 | - | - | - | - | - | - | - | - | - | - | - | - | - | - | - | - | -0.68** | - |
|  | 35 | - | - | - | - | - | - | - | - | - | - | - | - | - | - | - | - | - | -0.75** |
| *SD*: stomatal density (mm^-2^); *SS*: stomatal size (μm^2^); *a_max_*: maximum stomatal pore aperture (μm^2^); *ES*: Epidermal cell size (μm^2^); *a_op_*: operation stomatal aperture (μm^2^); *% a*: percentage of *a_op_* to *a_max_*; *g_max_*: maximum theoritical stomatal conductance (mol m^-2^ s^-1^); *IVD*: inter-veinal distance (mm); *VD*: total vein density (mm mm^-2^); *A_n_*: net carbon assimilation rate (μmol m^-2^ s^-1^); *g_s_*: stomatal conductance (mol m^-2^ s^-1^); i*WUE*: intrinsic water use efficiency (µmol CO_2_ mol^-1^ H_2_O); *LW*: leaf width (cm). | | | | | | | | | | | | | | | | | | | |

| **Table S7**. Pearson product-moment correlation analysis results for the relationships between the measured variables at different temperatures. The *R* coefficient and the statistical significance were determined using the mean value per line, per treatment for each variable. Statistical significance was judged as: *P*<0.001 (***), *P*<0.05 (**), *P*<0.1 (*), *P*>0.1 (ns). This table shows the relationships between leaf inner anatomy parameters and other anatomical and gas exchange parameters. (*n*=3) | | | | | | | | | | | | | | | | | | | | | | |
| --- | --- | --- | --- | --- | --- | --- | --- | --- | --- | --- | --- | --- | --- | --- | --- | --- | --- | --- | --- | --- | --- | --- |
|  | | *SD* | | | *SS* | | | *a_max_* | | | *ES* | | | *a_op_* | | | *% a* | | | *g_smax_* | | |
|  |  | 22 | 28 | 35 | 22 | 28 | 35 | 22 | 28 | 35 | 22 | 28 | 35 | 22 | 28 | 35 | 22 | 28 | 35 | 22 | 28 | 35 |
| *MT* | 22 | -0.66** | - | - | 0.57* | - | - | ns | - | - | ns | - | - | 0.79** | - | - | ns | - | - | ns | - | - |
|  | 28 | - | ns | - | - | ns | - | - | ns | - | - | 0.73* | - | - | ns | - | - | ns | - | - | ns | - |
|  | 35 | - | - | ns | - | - | 0.96*** | - | - | 0.9*** | - | - | 0.85** | - | - | ns | - | - | ns | - | - | ns |
| *IVD_c_* | 22 | ns | - | - | ns | - | - | ns | - | - | ns | - | - | 0.85** | - | - | 0.62* | - | - | ns | - | - |
|  | 28 | - | ns | - | - | ns | - | - | ns | - | - | ns | - | - | ns | - | - | ns | - | - | ns | - |
|  | 35 | - | - | ns | - | - | 0.63* | - | - | 0.7* | - | - | ns | - | - | ns | - | - | ns | - | - | ns |
| *M / BS* | 22 | ns | - | - | ns | - | - | ns | - | - | ns | - | - | ns | - | - | ns | - | - | ns | - | - |
|  | 28 | - | ns | - | - | ns | - | - | ns | - | - | ns | - | - | ns | - | - | ns | - | - | ns | - |
|  | 35 | - | - | ns | - | - | ns | - | - | ns | - | - | ns | - | - | ns | - | - | ns | - | - | ns |
| *MC_length_* | 22 | -0.66** | - | - | ns | - | - | ns | - | - | ns | - | - | 0.71** | - | - | 0.63** | - | - | -0.59* | - | - |
|  | 28 | - | ns | - | - | ns | - | - | ns | - | - | ns | - | - | ns | - | - | ns | - | - | ns | - |
|  | 35 | - | - | ns | - | - | 0.88** | - | - | 0.7* | - | - | ns | - | - | ns | - | - | ns | - | - | ns |
| *MC_area_* | 22 | -0.65** | - | - | ns | - | - | ns | - | - | 0.71* | - | - | 0.62* | - | - | ns | - | - | ns | - | - |
|  | 28 | - | -0.56* | - | - | ns | - | - | ns | - | - | 0.78** | - | - | ns | - | - | ns | - | - | -0.61* | - |
|  | 35 | - | - | -0.67* | - | - | 0.94*** | - | - | 0.76** | - | - | 0.8** | - | - | ns | - | - | ns | - | - | ns |
| *BSC_area_* | 22 | -0.62* | - | - | ns | - | - | ns | - | - | ns | - | - | 0.88*** | - | - | 0.58* | - | - | ns | - | - |
|  | 28 | - | ns | - | - | ns | - | - | ns | - | - | 0.7* | - | - | ns | - | - | 0.58* | - | - | ns | - |
|  | 35 | - | - | ns | - | - | 0.76** | - | - | 0.69* | - | - | 0.68* | - | - | ns | - | - | ns | - | - | ns |
| *M_si_* | 22 | -0.69** | - | - | ns | - | - | ns | - | - | ns | - | - | 0.56* | - | - | ns | - | - | ns | - | - |
|  | 28 | - | ns | - | - | ns | - | - | ns | - | - | ns | - | - | ns | - | - | ns | - | - | ns | - |
|  | 35 | - | - | ns | - | - | 0.66* | - | - | ns | - | - | ns | - | - | ns | - | - | ns | - | - | ns |
| *BS_si_* | 22 | -0.6* | - | - | 0.56* | - | - | ns | - | - | ns | - | - | 0.8** | - | - | ns | - | - | ns | - | - |
|  | 28 | - | ns | - | - | 0.72** | - | - | ns | - | - | ns | - | - | 0.7** | - | - | 0.79** | - | - | ns | - |
|  | 35 | - | - | ns | - | - | 0.78** | - | - | 0.76** | - | - | 0.69* | - | - | ns | - | - | ns | - | - | ns |
| *VB_si_* | 22 | ns | - | - | 0.6* | - | - | 0.68** | - | - | ns | - | - | ns | - | - | ns | - | - | ns | - | - |
|  | 28 | - | ns | - | - | 0.66* | - | - | ns | - | - | ns | - | - | ns | - | - | ns | - | - | ns | - |
|  | 35 | - | - | ns | - | - | ns | - | - | 0.74** | - | - | ns | - | - | ns | - | - | ns | - | - | ns |
| *IAS_si_* | 22 | ns | - | - | 0.58* | - | - | 0.63* | - | - | ns | - | - | 0.66* | - | - | ns | - | - | ns | - | - |
|  | 28 | - | ns | - | - | ns | - | - | ns | - | - | ns | - | - | ns | - | - | ns | - | - | ns | - |
|  | 35 | - | - | ns | - | - | 0.85** | - | - | 0.84** | - | - | 0.95*** | - | - | ns | - | - | ns | - | - | ns |
| *Dist_H_* | 22 | -0.71** | - | - | ns | - | - | ns | - | - | ns | - | - | 0.77** | - | - | ns | - | - | ns | - | - |
|  | 28 | - | ns | - | - | ns | - | - | ns | - | - | 0.85** | - | - | ns | - | - | ns | - | - | ns | - |
|  | 35 | - | - | -0.66* | - | - | 0.93*** | - | - | 0.9*** | - | - | 0.88*** | - | - | 0.65* | - | - | ns | - | - | ns |
| *MT*: leaf mesophyll thickness (μm); *IVD_c_*: interveinal Distance measured from leaf cross-sections (μm); *M : BS*: mesophyll to bundle sheath ratio; *MC_length_*: length of mesophyll cell (μm); *MC_area_*: area of single mesophyll cell (μm^2^); *BSC_area_*: area of single bundle sheath cell (μm^2^); *M_si_*: mesophyll surface area per *IVD_c_* (μm^2^ μm^-1^); *BS_si_*: bundle sheath surface area per *IVD_c_* (μm^2^ μm^-1^); *VB_si_*: vascular bundle surface area per *IVD_c_* (μm^2^ μm^-1^); *IAS_si_*: intercellular airspace surface area per *IVD_c_* (μm^2^ μm^-1^); *Dist_H_*: Hydraulic Distance (μm); *SD*: stomatal density (mm^-2^); *SS*: stomatal size (μm^2^); *a_max_*: maximum stomatal pore aperture (μm^2^); *ES*: Epidermal cell size (μm^2^); *a_op_*: operation stomatal aperture (μm^2^); *% a*: percentage of *a_op_* to *a_max_*; *g_max_*: maximum theoritical stomatal conductance (mol m^-2^ s^-1^); *IVD*: inter-veinal distance (mm); *DV*: total vein density (mm mm^-2^); *TLV*: Total Number of longitudinal veins; *LMA*: leaf mass per area (g m^-2^); *A_n_*: net carbon assimilation rate (μmol m^-2^ s^-1^); *g_s_*: stomatal conductance (mol m^-2^ s^-1^); i*WUE*: intrinsic water use efficiency (µmol CO_2_ mol^-1^ H_2_O); *LW*: leaf width (cm). | | | | | | | | | | | | | | | | | | | | | | |

| **Table 7 continued…** | | | | | | | | | | | | | | | | | | | | | | |
| --- | --- | --- | --- | --- | --- | --- | --- | --- | --- | --- | --- | --- | --- | --- | --- | --- | --- | --- | --- | --- | --- | --- |
|  | | *IVD* | | | *VD* | | | *TLV* | | | *LMA* | | | *A_n_* | | | *g_s_* | | | *iWUE* | | |
|  |  | 22 | 28 | 35 | 22 | 28 | 35 | 22 | 28 | 35 | 22 | 28 | 35 | 22 | 28 | 35 | 22 | 28 | 35 | 22 | 2 8 | 35 |
| *MT* | 22 | 0.61* | - | - | -0.7** | - | - | ns | - | - | ns | - | - | ns | - | - | ns | - | - | ns | - | - |
|  | 28 | - | ns | - | - | ns | - | - | 0.61* | - | - | ns | - | - | ns | - | - | ns | - | - | ns | - |
|  | 35 | - | - | 0.61* | - | - | ns | - | - | 0.85** | - | - | ns | - | - | ns | - | - | ns | - | - | -0.74** |
| *IVD_c_* | 22 | 0.66** | - | - | -0.7** | - | - | ns | - | - | ns | - | - | ns | - | - | 0.58* | - | - | -0.78** | - | - |
|  | 28 | - | 0.89** | - | - | ns | - | - | ns | - | - | ns | - | - | ns | - | - | ns | - | - | ns | - |
|  | 35 | - | - | 0.83** | - | - | -0.8** | - | - | ns | - | - | ns | - | - | ns | - | - | ns | - | - | -0.86** |
| *M / BS* | 22 | -0.62* | - | - | ns | - | - | ns | - | - | -0.65** | - | - | ns | - | - | ns | - | - | ns | - | - |
|  | 28 | - | ns | - | - | ns | - | - | ns | - | - | ns | - | - | ns | - | - | ns | - | - | ns | - |
|  | 35 | - | - | ns | - | - | ns | - | - | ns | - | - | ns | - | - | ns | - | - | ns | - | - | ns |
| *MC_length_* | 22 | ns | - | - | -0.7** | - | - | ns | - | - | ns | - | - | ns | - | - | ns | - | - | -0.55* | - | - |
|  | 28 | - | 0.76** | - | - | -0.72** | - | - | ns | - | - | ns | - | - | ns | - | - | ns | - | - | ns | - |
|  | 35 | - | - | ns | - | - | ns | - | - | 0.76** | - | - | ns | - | - | ns | - | - | ns | - | - | ns |
| *MC_area_* | 22 | ns | - | - | -0.58* | - | - | ns | - | - | ns | - | - | ns | - | - | ns | - | - | ns | - | - |
|  | 28 | - | 0.7* | - | - | ns | - | - | ns | - | - | ns | - | - | ns | - | - | ns | - | - | ns | - |
|  | 35 | - | - | ns | - | - | ns | - | - | 0.8** | - | - | ns | - | - | ns | - | - | ns | - | - | -0.6* |
| *BSC_area_* | 22 | 0.83** | - | - | -0.83** | - | - | ns | - | - | ns | - | - | ns | - | - | ns | - | - | -0.71** | - | - |
|  | 28 | - | 0.85** | - | - | ns | - | - | ns | - | - | ns | - | - | ns | - | - | ns | - | - | ns | - |
|  | 35 | - | - | 0.63* | - | - | ns | - | - | ns | - | - | ns | - | - | ns | - | - | ns | - | - | -0.63* |
| *M_si_* | 22 | ns | - | - | ns | - | - | ns | - | - | ns | - | - | ns | - | - | ns | - | - | ns | - | - |
|  | 28 | - | ns | - | - | ns | - | - | ns | - | - | ns | - | - | ns | - | - | ns | - | - | ns | - |
|  | 35 | - | - | ns | - | - | ns | - | - | 0.67* | - | - | ns | - | - | ns | - | - | ns | - | - | ns |
| *BS_si_* | 22 | 0.74** | - | - | -0.81** | - | - | ns | - | - | ns | - | - | ns | - | - | ns | - | - | -0.57* | - | - |
|  | 28 | - | ns | - | - | ns | - | - | 0.58* | - | - | ns | - | - | ns | - | - | 0.64** | - | - | ns | - |
|  | 35 | - | - | 0.63* | - | - | ns | - | - | 0.67* | - | - | ns | - | - | ns | - | - | ns | - | - | -0.63* |
| *VB_si_* | 22 | ns | - | - | ns | - | - | 0.67** | - | - | ns | - | - | ns | - | - | ns | - | - | ns | - | - |
|  | 28 | - | ns | - | - | ns | - | - | ns | - | - | ns | - | - | ns | - | - | ns | - | - | ns | - |
|  | 35 | - | - | 0.63* | - | - | ns | - | - | ns | - | - | ns | - | - | ns | - | - | ns | - | - | ns |
| *IAS_si_* | 22 | 0.63* | - | - | ns | - | - | ns | - | - | 0.85** | - | - | ns | - | - | ns | - | - | ns | - | - |
|  | 28 | - | ns | - | - | ns | - | - | ns | - | - | ns | - | - | ns | - | - | ns | - | - | ns | - |
|  | 35 | - | - | ns | - | - | ns | - | - | 0.68* | - | - | 0.59* | - | - | ns | - | - | ns | - | - | -0.73** |
|  | 22 | 0.62* | - | - | -0.74** | - | - | ns | - | - | ns | - | - | ns | - | - | ns | - | - | ns | - | - |
| *Dist_H_* | 28 | - | ns | - | - | ns | - | - | ns | - | - | ns | - | - | ns | - | - | ns | - | - | ns | - |
|  | 35 | - | - | 0.67* | - | - | -0.63* | - | - | 0.8** | - | - | ns | - | - | ns | - | - | ns | - | - | -0.82** |
| *MT*: leaf mesophyll thickness (μm); *IVD_c_*: interveinal Distance measured from leaf cross-sections (μm); *M : BS*: mesophyll to bundle sheath ratio; *MC_length_*: length of mesophyll cell (μm); *MC_area_*: area of single mesophyll cell (μm^2^); *BSC_area_*: area of single bundle sheath cell (μm^2^); *M_si_*: mesophyll surface area per *IVD_c_* (μm^2^ μm^-1^); *BS_si_*: bundle sheath surface area per *IVD_c_* (μm^2^ μm^-1^); *VB_si_*: vascular bundle surface area per *IVD_c_* (μm^2^ μm^-1^); *IAS_si_*: intercellular airspace surface area per *IVD_c_* (μm^2^ μm^-1^); *Dist_H_*: Hydraulic Distance (μm); *SD*: stomatal density (mm^-2^); *SS*: stomatal size (μm^2^); *a_max_*: maximum stomatal pore aperture (μm^2^); *ES*: Epidermal cell size (μm^2^); *a_op_*: operation stomatal aperture (μm^2^); *% a*: percentage of *a_op_* to *a_max_*; *g_max_*: maximum theoritical stomatal conductance (mol m^-2^ s^-1^); *IVD*: inter-veinal distance (mm); *DV*: total vein density (mm mm^-2^); *TLV*: Total Number of longitudinal veins; *LMA*: leaf mass per area (g m^-2^); *A_n_*: net carbon assimilation rate (μmol m^-2^ s^-1^); *g_s_*: stomatal conductance (mol m^-2^ s^-1^); i*WUE*: intrinsic water use efficiency (µmol CO_2_ mol^-1^ H_2_O); *LW*: leaf width (cm). | | | | | | | | | | | | | | | | | | | | | | |

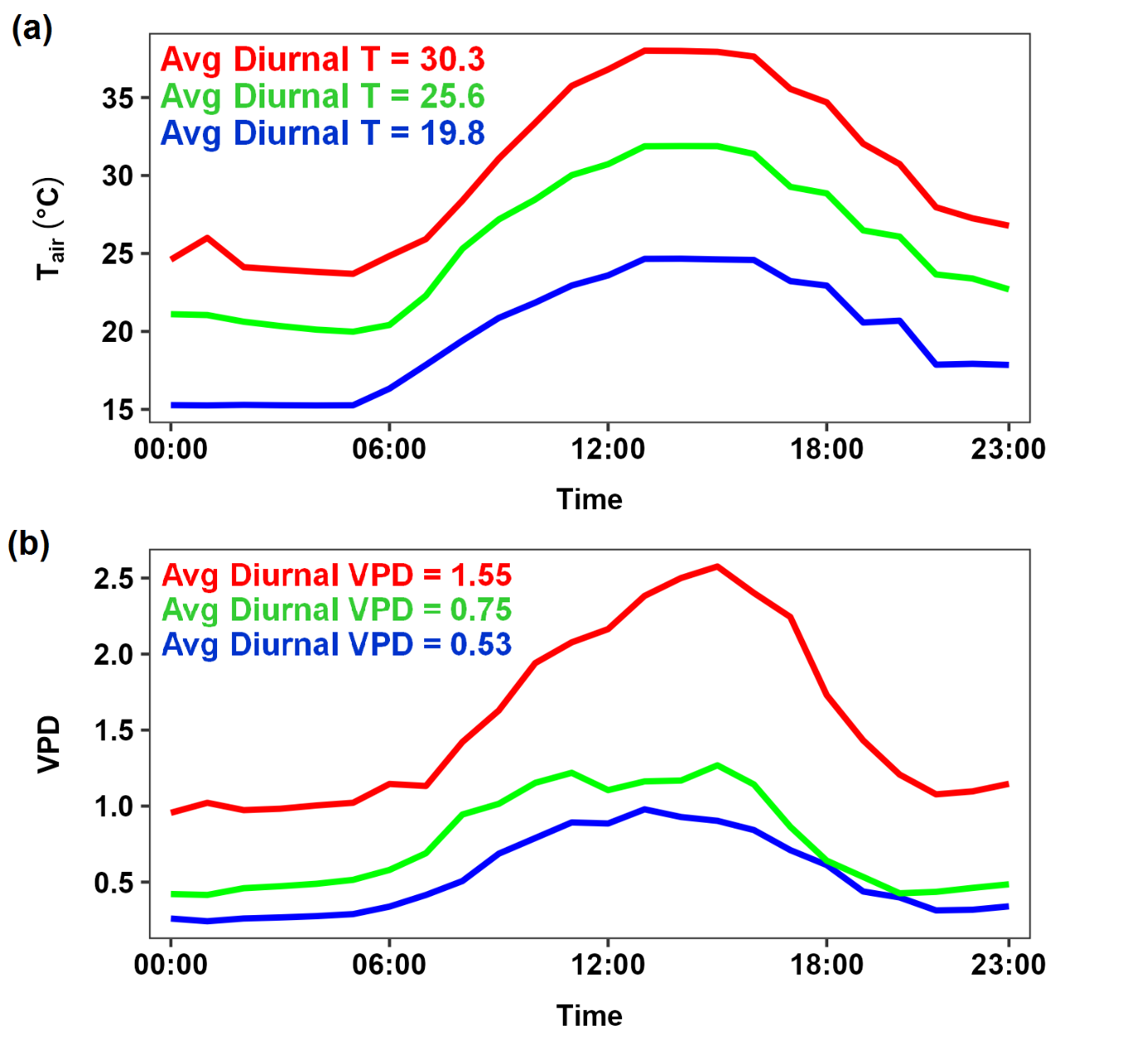

**Fig. S1** Plots showing the average environmental conditions recorded in the glasshouse chambers over a day. The plots are the average of ALL the growth and measurement period per hour (14/10/2016 to 26/11/2016). High temperature chamber is in red, mid-temperature treatment in green and low-temperature treatment in blue.

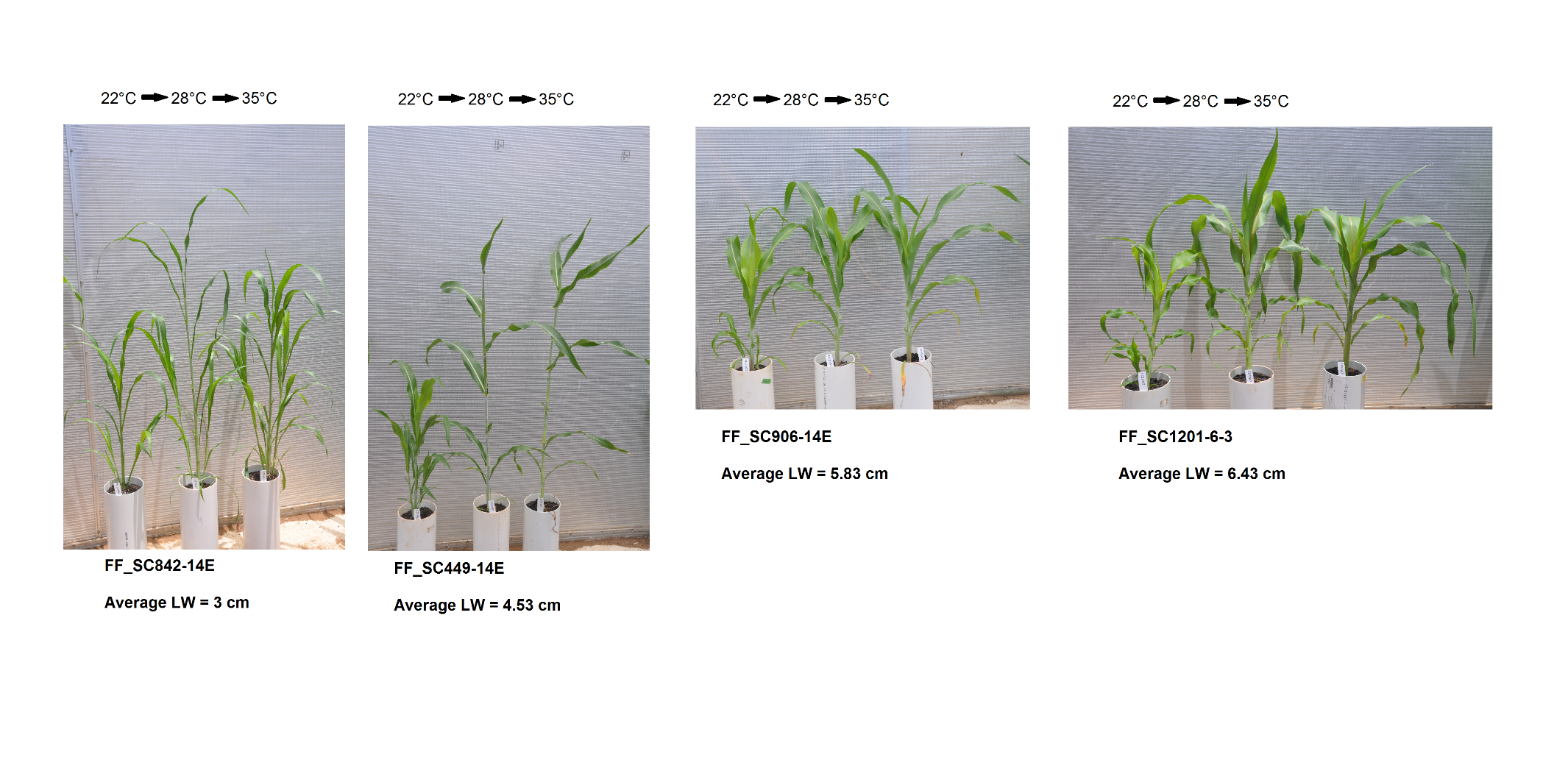

**Fig. S2**	A selection of plants representing different lines of different average leaf width (*LW*). Average *LW* is the mean of all the replicates per lines with treatments combine. The images were taken around the time of measurement (46 days after emergence).

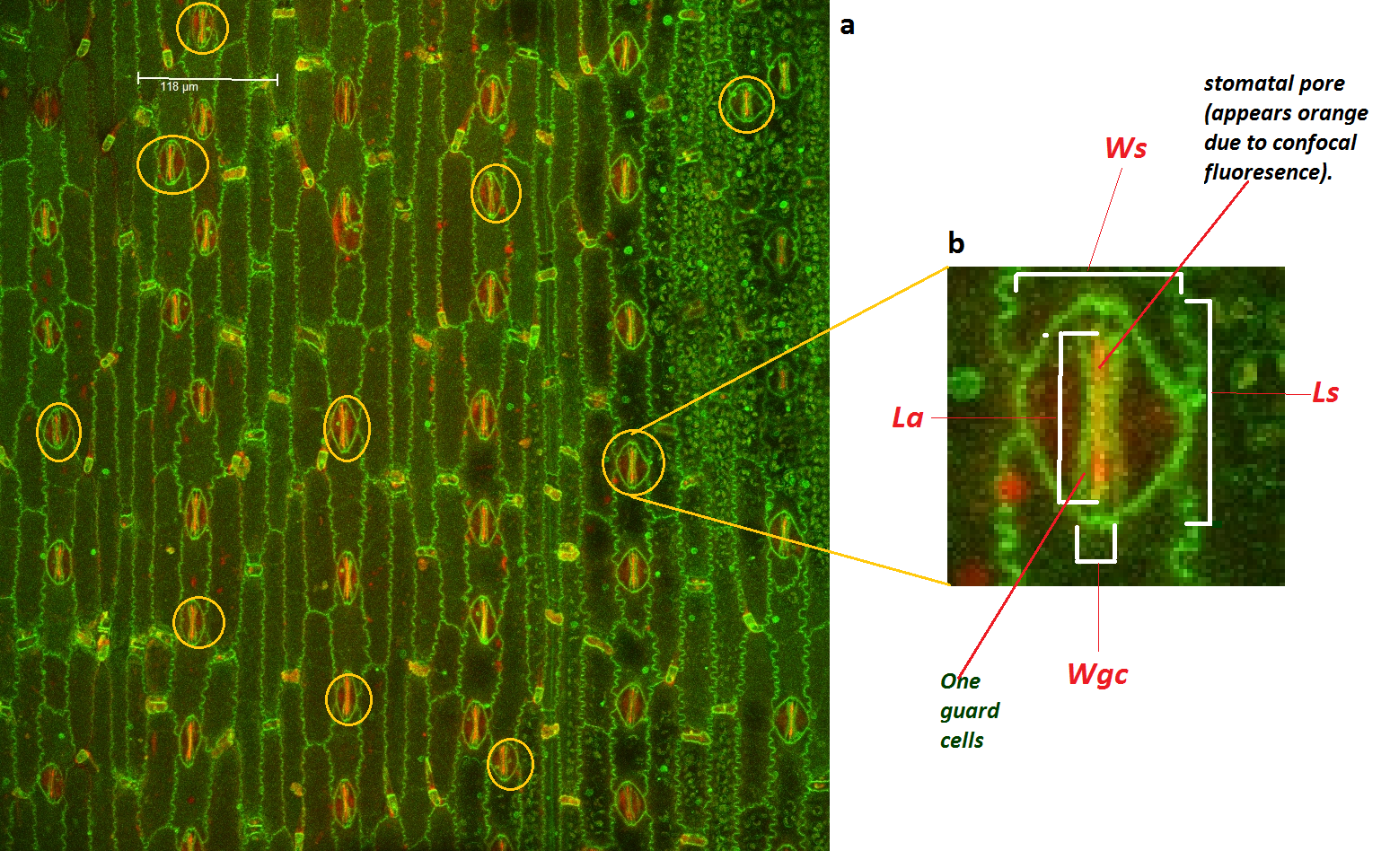
 **Fig. S3** Confocal images to illustrate sampling of stomatal features. **(a)** shows **all** the area used for *SD* measurements with scale, and an example of the 10 randomly selected stoma identified for *SS* measurement. **(b)** shows stomatal dimensions: Width of stomata complex (*W_s_*) (with subsidiary cells), width of stomatal guard cells (*W_gc_*), length of stomata (*L_s_*), and length of aperture (*L_a_*).

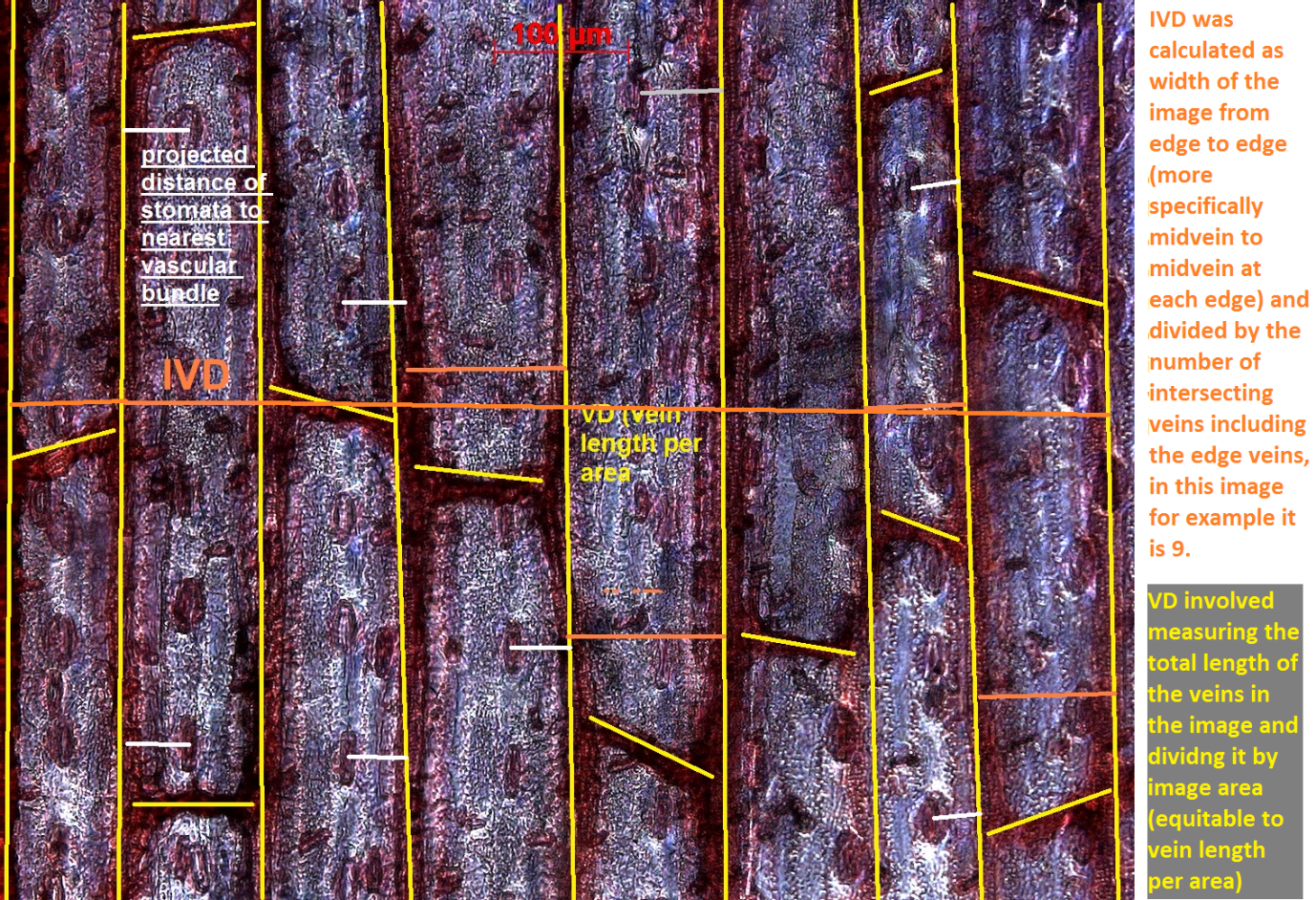

**Fig. S4** Light microscopy image of a cleared and stained leaf showing the veins in yellow lines. Full length of the yellow lines per area compromised vein density (*VD –* or vein length per area). interveinal distance (*IVD*) is also highlighted in orange (middle of vascular bundle to middle of vascular bundle). The projected distance between the stomata and middle of the nearest vascular bundle was used in the calculation of hydraulic distance (*Dist_H_*).

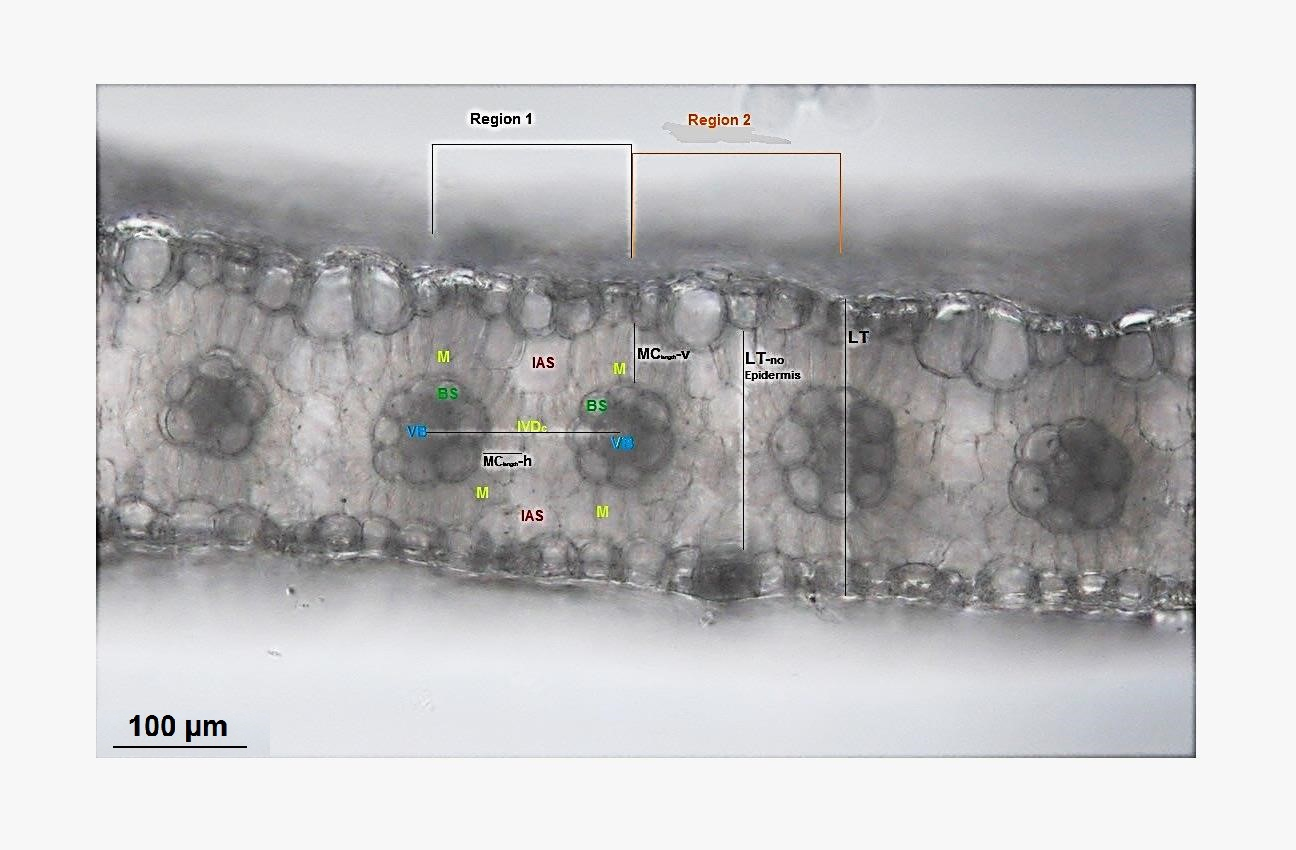

**Fig. S5** Light microscopy image of a cleared transverse leaf cross-section showing the main anatomical components that were measured. IAS: Intercellular air spaces, M: Mesophyll cells; BS: Bundle sheath cells; VB: Vascular bundle; *IVD_c_*: Interveinal distance; *LT*: Leaf Thickness; *LT-*no epidermis: Leaf Mesophyll Thickness; *MC_length_*-h: length of horizontal mesophyll cells; *MC_length_*-v: length of vertical mesophyll cells.

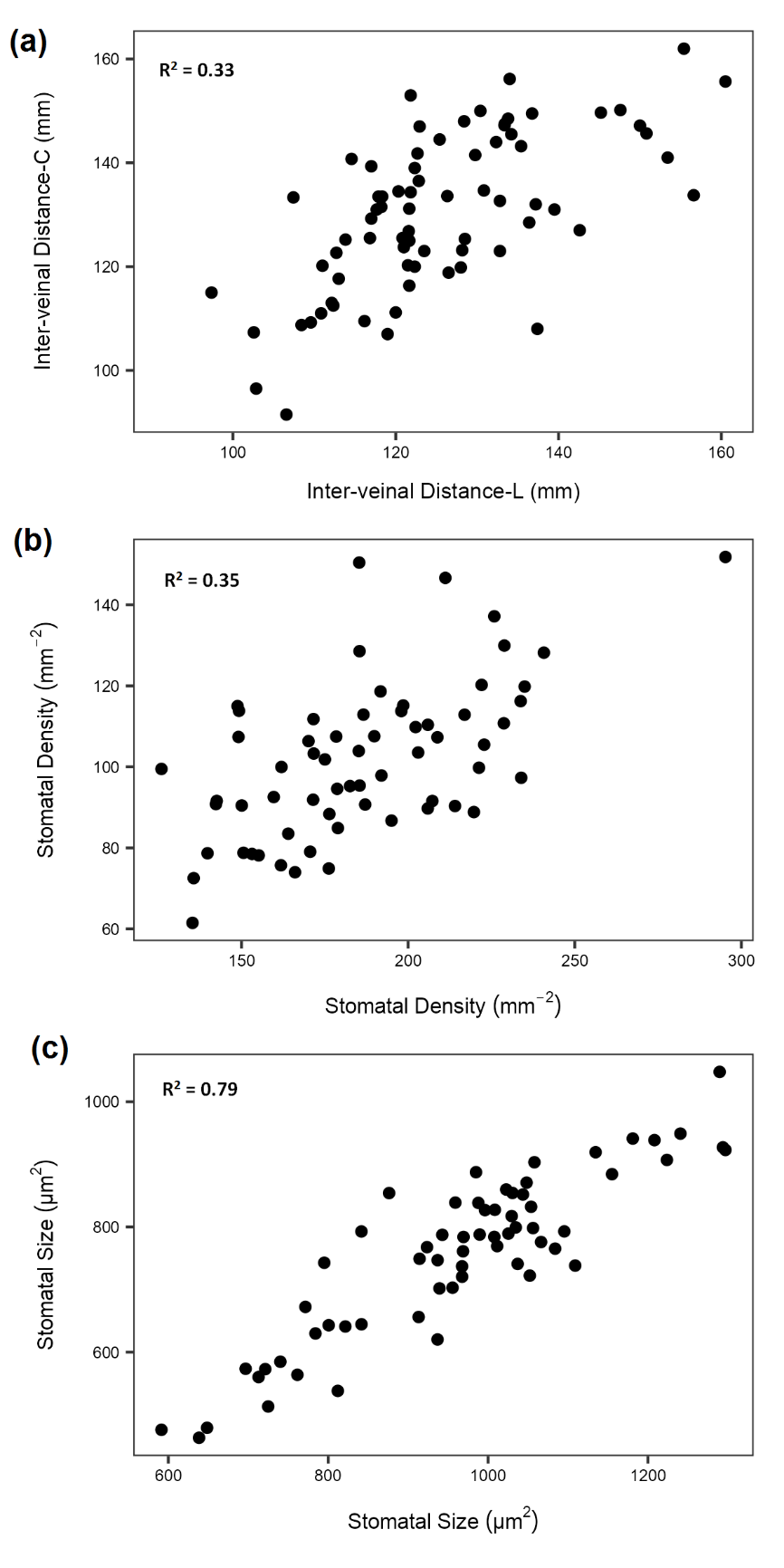

**Fig. S6.** Plots of data points of (a) Interveinal distance measured from cross-sections (y-axis) vs. Interveinal distance measured from paradermal sections (x-axis). (b) Stomatal density measured during first stage of analysis (x-axis) vs. Stomatal density measured later when analysing for epidermal cell size (y-axis). (c) Similar to (b) but for stomatal size.

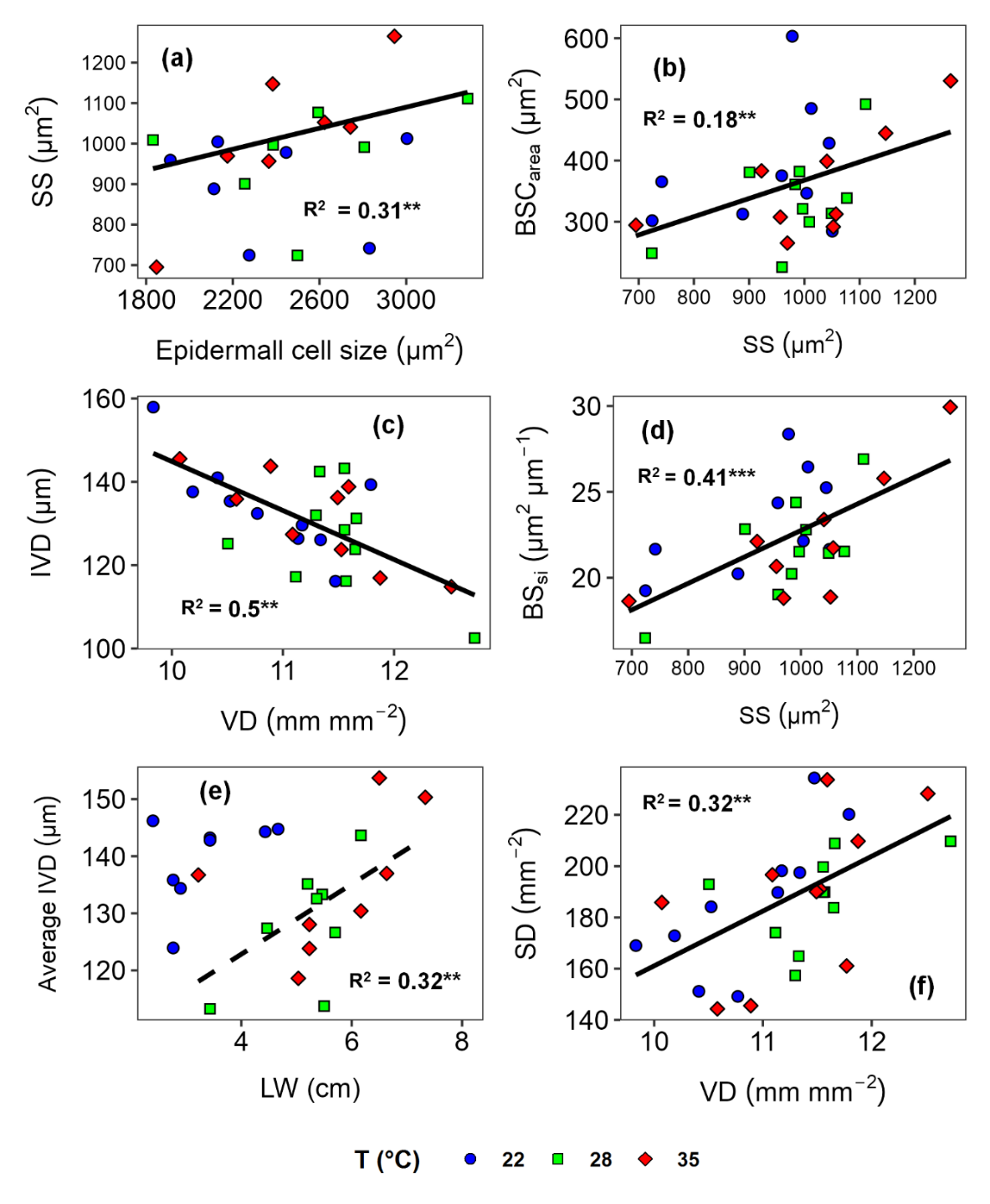

**Fig.** **S7** Relationships between stomatal size, leaf width and leaf anatomy in ten *Sorghum bicolor* lines grown under different temperatures. Leaf width was measured on the portion sampled from the middle of the youngest fully expanded leaf. The same portion was used for measurement of vein and stomatal traits by analysing the area between 2nd and 3rd major veins from the midrib. Each point represents the mean value per line per treatment (*n*=3). Standard error bars were removed to ensure clearer presentation (Tables S1, S2 and S3). Pearson correlation analyses were conducted and significant results were shown with a solid line with the corresponding *R^2^* value (*P*<0.001=***, *P*<0.05=**, *P*<0.1=*). Dashed black line indicate a significant correlation when the 22°C treatment was removed (Table S5). Growth temperatures were: blue=22°C, green=28°C, red=35°C. (a) Stomatal size (*SS*) vs Epidermal cell size; (b) *SS* vs. Bundle sheath cell area (*BSC_area_*).; (c) Inter-veinal distance (*IVD*) vs Vein density (*VD*); (d) Cross-sectional bundle sheath surface area per *IVD* (*BS_si_*) vs *SS*; (e) *IVD* vs. *LW*; (f) *SD* vs. *VD*. **NB**: for Fig. (e), each area between two **major** veins was taken and the *IVD* between **minor** veins of that specific area was calculated. The *IVD* values of the different leaf segments along the leaf width were pooled and the mean is presented in Fig. (e) as “Average *IVD*”.

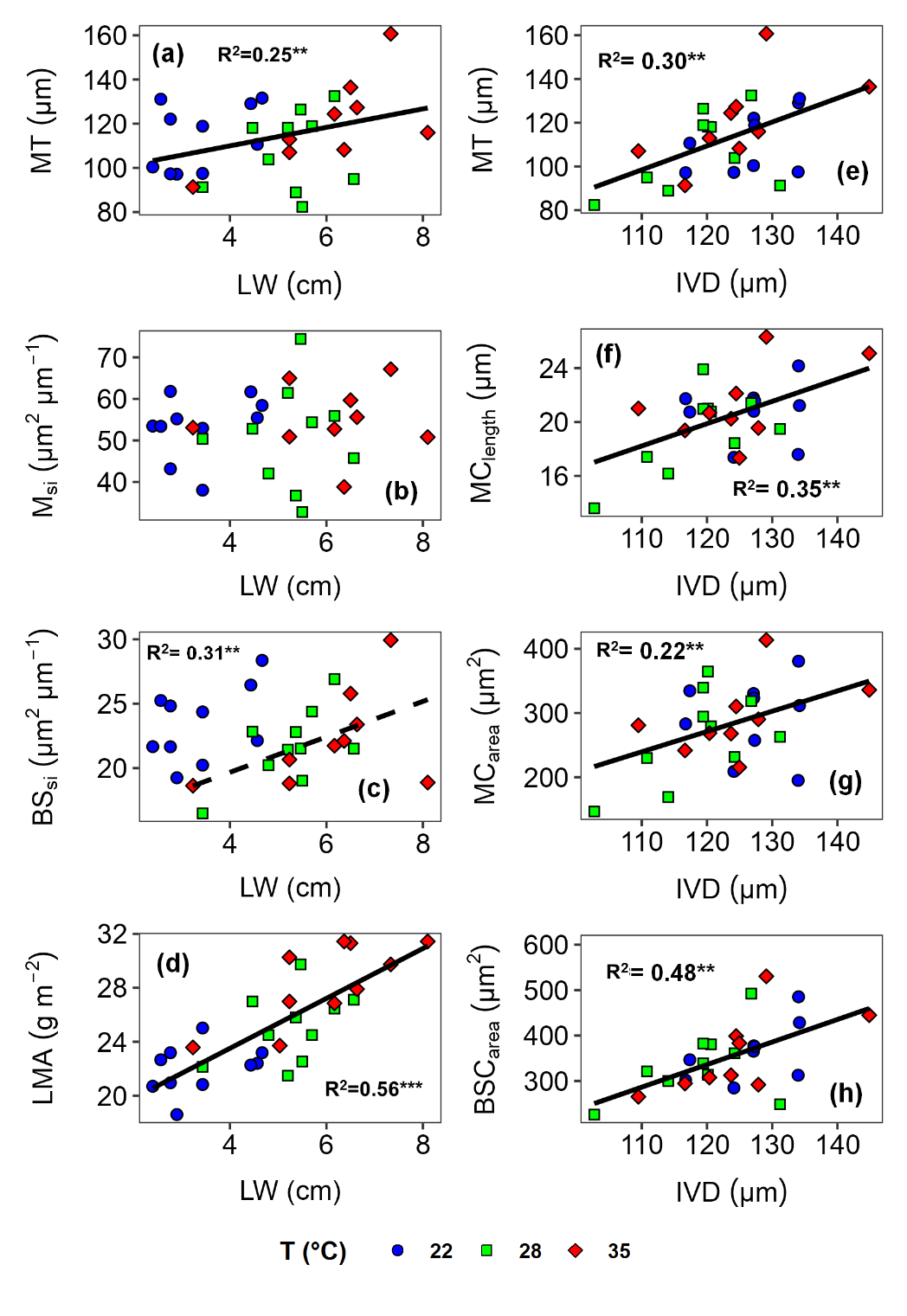

**Fig.** **S8** Relationships between leaf width (*LW*), interveinal distance (*IVD*) and inner leaf anatomy in ten *Sorghum bicolor* lines grown under different temperatures. Leaf width was measured on the portion sampled from the middle of the youngest fully expanded leaf. The same portion was used for measurement of vein and inner anatomy traits by analysing the area between 2nd and 3rd major veins. Each point represents the mean per line per treatment (n=3). Standard error bars were removed to ensure clearer presentation (Tables S1, S3 and S4). Pearson correlation analyses were conducted and significant results were shown with a solid line with the corresponding *r^2^* value (*P*<0.001=***, *P*<0.05=**, *P*<0.1=*). Dashed black line indicate a significant correlation when the 22°C treatment was removed (Table S5). Different growth temperatures are represented by the different fill colour of the scatter points: blue=22°C, green=28°C, red=35°C. *LW* vs. (a) Leaf mesophyll thickness (*MT*); (b) Cross-sectional mesophyll surface area per *IVD* (*M_si_*); (c) Cross-sectional bundle sheath surface area per *IVD* (*BS_si_*); (d) Leaf mass per area (*LMA*); *IVD* vs. (e) *MT*; (f) Mesophyll cell length (*MC_length_*) (g) Mesophyll cell area (*MC_area_*); (h) Bundle sheath cell area (*BSC_area_*).

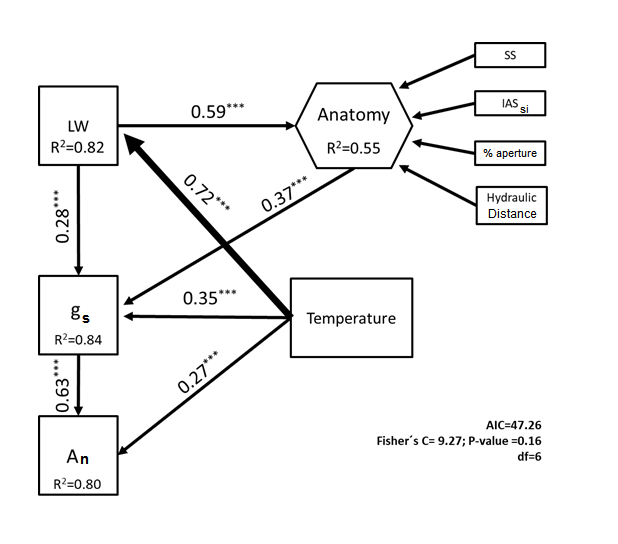

Fig. S9 Effects of growth temperature on leaf width, leaf anatomy, stomatal conductance (*g_s_*) and carbon assimilation rate (*A_n_*) across genotypes. Numbers adjacent to the arrows indicate the effect size of the relationship, while asterisks denote *P*-values at <0.0001. Leaf anatomy was included as a composite variable, and included the following variables: stomatal size (*SS*), Cross-sectional intercellular airspace surface area per inter-veinal distance (*IAS_si_*), operational stomatal aperture as a percentage of the maximum *(% aperture*) and hydraulic distance.  *R^2^* denotes the proportion of the model variance explained. Overall goodness-of-fit test is shown in the bottom of the figure.
